# Integrated histopathology of the human pancreas throughout stages of type 1 diabetes progression

**DOI:** 10.1101/2025.03.18.644000

**Authors:** Verena van der Heide, Sara McArdle, Michael S. Nelson, Karen Cerosaletti, Sacha Gnjatic, Zbigniew Mikulski, Amanda L. Posgai, Irina Kusmartseva, Mark Atkinson, Dirk Homann

**Affiliations:** Marc and Jennifer Lipschultz Precision Immunology Institute, Department of Immunology and Immunotherapy, Icahn School of Medicine at Mount Sinai (ISMMS), New York, NY 10029, USA; Microscopy and Histology Core Facility, La Jolla Institute for Immunology, La Jolla, CA, 92037, USA; Department of Biomedical Engineering, University of Wisconsin-Madison, Madison, WI 53706, USA; Center for Translational Immunology, Benaroya Research Institute, Seattle, WA 98101, USA; Tisch Cancer Institute, Department of Medicine, ISMMS, New York, NY 10029, USA; Department of Pathology, Immunology, and Laboratory Medicine, University of Florida Diabetes Institute, College of Medicine, Gainesville, FL 32610, USA; Department of Pediatrics, University of Florida Diabetes Institute, College of Medicine, Gainesville, FL 32610, USA; Diabetes, Obesity & Metabolism Institute, Department of Medicine, ISMMS, New York, NY 10029, USA

**Keywords:** type 1 diabetes (T1D), endocrine pancreas, islet of Langerhans, beta cells, immune cells, histopathology, multiplex imaging, whole slide imaging, digital pathology, QuPath, semi-automated image analysis

## Abstract

Type 1 diabetes (T1D) is a progressive autoimmune condition that culminates in the loss of insulin-producing beta cells. Pancreatic histopathology provides essential insights into disease initiation and progression yet an integrated perspective of *in situ* pathogenic processes is lacking due to limited sample availability, the dispersed nature of anatomical lesions, and often restricted analytical dimensionality. Here, we combined multiplexed immunostaining, high-magnification whole-slide imaging, digital pathology, and semi-automated image analysis strategies to interrogate pancreatic tail and head regions obtained from organ donors across T1D stages including at-risk and at-onset cases. Deconvolution of architectural features, endocrine cell composition, immune cell burden, and spatial relations of ∼25,000 islets revealed a series of novel histopathological correlates especially in the prodromal disease stage preceding clinical T1D. Altogether, our comprehensive “single-islet” analyses permit the reconstruction of a revised natural T1D history with implications for further histopathological investigations, considerations of pathogenetic modalities, and therapeutic interventions.

## INTRODUCTION

Type 1 diabetes (T1D) is a chronic autoimmune disease characterized by the loss of insulin-producing beta cells from pancreatic islets^1,2^. An interplay of genetic susceptibility^3^, environmental factors^4^, and immune dysregulation^5^ promotes multiple pathological alterations including focal lesions in the islets (insulitis), pancreatic inflammation, reduced exocrine pancreas volume, and systemic complications due to overt hyperglycemia^6,7^. Early histological interrogations of the pancreas established T1D as an autoimmune disease^8^ and defined pathological hallmarks for the organ^9^, yet extensive *in situ* studies have mostly been limited to the past ∼15 years following the creation of specialized tissue repositories such as the Network for Pancreatic Organ donors with Diabetes (nPOD) program^10^. Nevertheless, estimates suggest that <700 T1D donor pancreata are available for research globally^11^, a limited resource that is further compounded by substantial heterogeneity at the level of organ donors, pancreas anatomy (*e.g.,* organ region, lobular insulitis), islet properties (*e.g.,* architecture, endocrine composition, association with immune cells), and altered endocrine function beyond beta cells (*e.g.,* alpha cells^12,13^); pancreata from donors at high risk (≥2 auto-antibodies [AAbs], elevated HLA class-II risk) and at-onset of clinical T1D, essential for pathogenesis studies, are exceptionally scarce.

To contend with these challenges, emerging investigative techniques such as whole-slide imaging^9,14–25^, high-dimensional tissue imaging^24,26–31^, and three-dimensional (3D) visualization of immuno-stained thick pancreas slices^32,33^ have been deployed in conjunction with tailored digital analysis software. However, to date no standardized data analytics platform dedicated to pancreatic histopathology is readily accessible. The commercial HALO platform, featuring “pancreas modules” and machine learning algorithms, has aided the identification of pancreas and islet features^17,19,24,29,34–36^, but its proprietary nature restricts customization. QuPath, a versatile open-source digital pathology platform, is gaining popularity^37,38^ and has been employed in recent T1D studies primarily focused on immune cell analyses^20–23,39^.

To our knowledge, an integration of the complementary strands of histopathological T1D pancreas investigation and digital pathology modalities^10,40^ has not yet been performed. We therefore combined higher dimensional multiplex brightfield immunohistochemistry (IHC), whole-slide image acquisition at high magnification, and a novel semi-automated digital analysis pipeline implemented in QuPath to trace the histopathology of T1D development and progression. Our results reveal a spatially homogenous and islet size-contingent architectural organization of the endocrine pancreas, a notable coordination of organ-wide pathogenic processes supported by an apparent absence of a simple correlation between islet composition and associated immune cell burden, and multiple distinct histopathological correlates that foreshadow T1D histopathology already at the preclinical stage. Accordingly, we propose a revised natural history of T1D that effectively contextualizes established hallmarks of T1D pathology with multiple novel observations.

## RESULTS & DISCUSSION

### Multiplexed IHC analysis of human pancreatic tissue sections

The detailed histopathological interrogation of healthy and diabetic human pancreatic sections remains a cornerstone of investigations into T1D pathogenesis^10^. To leverage recent advances in tissue imaging, we employed the Multiplexed Immunohistochemical Consecutive Staining on Single Slide (MICSSS) platform^41,42^, an iterative IHC assay that permits successive brightfield visualization of immuno-staining patterns at high magnification (40x) across whole slides (***Figs.1A/B***). Previous MICSSS applications were largely restricted to characterization of CD45^+^ hematopoietic cells^43–45^, and our adaptation for the visualization of eight pancreatic endocrine hormones in addition to CD45, in part due to larger intracellular staining areas, required adjustments to signal amplification, destaining/blocking, and an empirical determination of staining order since our targets of interest exhibited differential sensitivity to deterioration over repeated tissue processing steps (***Figs.1C***).

### Development of a semi-automated image analysis pipeline for MICSSS-stained pancreatic tissue sections

We used machine learning to analyze pancreatic whole-slide images and customized a series of specific tasks in the open-source digital pathology software QuPath^37^. We first captured total tissue specimen areas and excluded surrounding connective tissue and fat to demarcate parenchymal (exocrine and endocrine) areas (***Fig.1D***). Alignment of nine immuno-stained images per tissue section was performed using affine transformation and achieved cell-level accuracy. Pancreatic islets, defined here as endocrine cell clusters ≥1,000μm^2^ (∼10 cells/∼36μm diameter), were delineated by merging the areas of six selected hormone stains with residual non-endocrine areas to create contiguous objects (***Fig.1E*** and Methods). Geometric and spatial properties of islet outlines were quantified to allow for a basic description of islet architecture and derivation of shape descriptors (***Fig.1F***).

**Figure 1.**
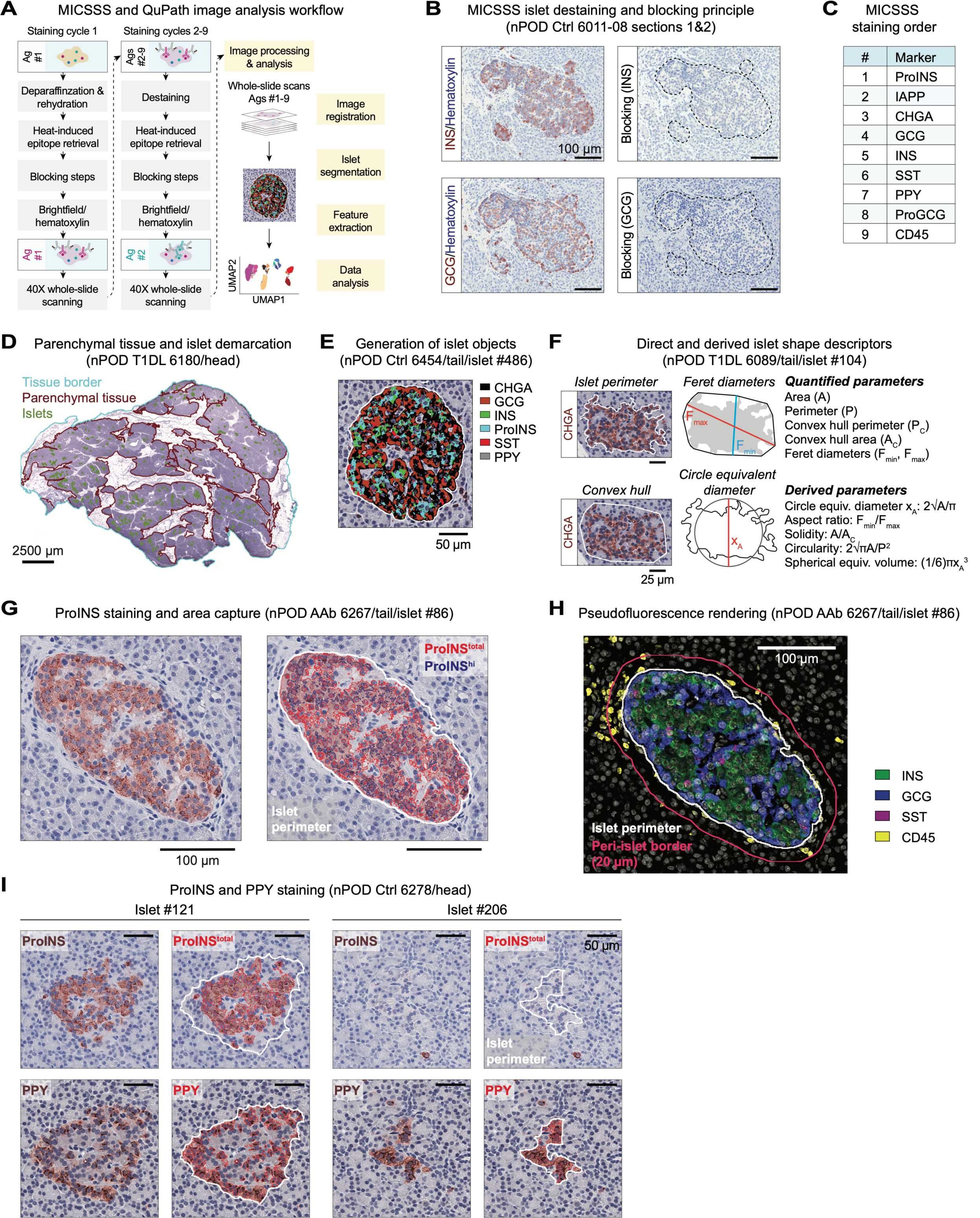
MICSSS staining and semi-automated image analysis of pancreatic tissue sections. **A.**, workflow of MICSSS staining and subsequent image analysis pipeline. In subsequent panels B.-I., donor type (Ctrl: non-diabetic, AAb: stage 1/2 T1D, T1DL: longer duration T1D), nPOD ID and pancreatic region are indicated. **B.,** MICSSS destaining/blocking illustrated for INS and GCG stains on adjacent slides. **C.,** MICSSS staining order. **D.,** demarcation of total tissue, parenchymal (exocrine/endocrine) tissue and islet areas. **E.,** generation of islet objects by additive display of six endocrine hormone stains (no PPY in present islet). **F.,** islet shape measurements and equations for derived parameters. **G.,** ProINS brightfield stain (brown), automated capture of ProINS staining areas (ProINS^total^: red traces, ProINS^hi^: blue traces), and islet perimeter (white). **H.,** pseudo-fluorescent rendering of overlaid INS, GCG, SST and CD45 stains, and demarcation of peri-islet region (border at 20μm distance from islet perimeter). **I.,** ProINS and PPY staining (brown) and respective area captures (red traces) for representative ProINS^+^PPY^+^ and ProINS^-^PPY^+^ islets.

To delimit hormone staining areas within each islet, islet outlines were superimposed onto each immuno-stained image and a pixel classifier was used to segment positive staining areas. Given the complete lack of background signal and heterogeneous pattern of proinsulin (ProINS) expression as observed previously (high: Golgi/immature INS granules; low: cytoplasm/ER)^46,47^, we quantified islet areas of both high and total ProINS content (***Fig.1G***). Representative images for other hormone staining area captures are shown in ***Fig.S1A***, and due to the creeping introduction of some background signal over successive staining rounds, we refrained from distinguishing areas of low and high expression for all other hormones thus permitting proper demarcation of mutually exclusive staining areas (***Fig.1H***). The segmented regions from each hormone stain were further used to calculate total endocrine areas (*i.e.,* union of eight hormone stains) and Jaccard indices, a statistic that quantifies the overlap of brightfield staining areas and corroborates the accuracy of our image alignment/segmentation strategy (***Fig.S1B***). Lastly, we incorporated the StarDist algorithm^48^ and an object classifier into our analysis pipeline to capture CD45^+^ hematopoietic cells within islets and the surrounding peri-islet areas (***Figs.1H & S1A***).

The importance for the present study to include the visualization of pancreatic polypeptide (PPY) expression, largely segregated to the uncinate process of the pancreatic head (PH)^49–51^, is illustrated by our observation that some PPY^+^ islets lack alpha and beta cells even in non-diabetic donors (***Figs.1I & S1C***); thus, any consideration of insulin-deficient islets (IDIs) in the PH needs to include PPY analyses. Altogether, our automation of multiple image analysis tasks in combination with multiplexed whole-slide image acquisition offers unique opportunities to revisit pancreatic histology in health and T1D disease *at scale*.

### Pancreatic donor cohorts and basic endocrine hormone expression patterns across T1D stages

To interrogate the histopathological evolution of T1D, we performed MICSSS staining of pancreatic tail (PT) and PH sections from four donor groups: non-diabetic controls (Ctrl), auto-antibody-positive stage 1/2 T1D (AAb), short duration stage 3 T1D (T1DS; <2 years including three at-onset donors), and longer duration T1D (T1DL; 8-11 years) (***Tables 1 & S1***). Donor matching was performed on age and gender and, where possible, demographic (ethnicity) and clinical (body mass index/BMI) parameters; several outliers in our cohorts are described in ***Table S1***. HLA-II risk^52–55^ was calculated to be low for Ctrl subjects and similarly elevated for AAb, T1DS and T1DL donors (***Table S1 & Fig.S2A***). Age, BMI, and total pancreas weight display a significant positive correlation for all donors as expected^56^; in T1DS and T1DL groups, donor age and age of T1D onset correlate near perfectly; and C-peptide and HbA1c exhibit a good fit in a one-phase exponential decay model (***Fig.S2B/C***).

Consistent with the T1D-associated loss of pancreas mass^56,57^, relative pancreas weights tend to decline in T1DS subjects, with a somewhat more pronounced reduction in our T1DL cohort (***Fig.S2D***). Interestingly, this decrease is also reflected in a trend toward reduction of absolute and relative parenchymal tissue section areas, and PT but not PH parenchymal areas further correlate positively with age (***Fig.S2E/F***). On average, we captured ∼500 islets per tissue section, with absolute numbers broadly decreasing as T1D progresses (***Fig.S2G***). While islet densities show a similar 1.5-2.0-fold non-significant reduction across donor cohorts, cumulative islet areas decline significantly (Ctrl *vs.* T1DL PH: 1.8-fold; PT: 2.9-fold) corresponding to an overall 2.9-5.1-fold loss of islet mass (***Figs.2A & S2H***). Quantifying cumulative islet hormone staining areas as a fraction of parenchymal tissue areas provides an initial orientation. As in earlier whole-slide imaging studies^14,15^, insulin (INS) staining areas are significantly reduced in T1DS/T1DL donors while glucagon (GCG) areas remain largely preserved (***Fig.S2I***). Several additional observations are noteworthy: a ∼2.7-fold loss of chromogranin A (CHGA) from Ctrl to T1DL stage accompanies the decline of relative islet areas; total ProINS areas are larger than INS areas whereas ProINS^hi^ areas are smaller and comparable to a previous report^15^; a significant decrease of ProINS and islet amyloid polypeptide (IAPP) areas in PH but not PT of AAb donors strikingly echoes recent findings about lower beta cell volume assessed by 3D morphometric analyses^33^; somatostatin (SST) staining areas remain unaltered in all donor groups; and PPY expression, minor in PT but prominent in PH, also appears unaffected by disease progression (***Fig.S2I***).

For additional context, we document a progressive decline of INS:GCG expression ratios (***Fig.S2J***) as well as an estimation of total endocrine cell type mass. Beta cell mass wanes precipitously (Ctrl *vs.* T1DL ProINS: ∼780-fold; INS: ∼1,250-fold), but both alpha (ProGCG/GCG) and delta (SST) cell mass also decrease in T1DL donors, albeit to a much lesser degree (∼3-fold; PT>PH). In contrast, gamma cell (PPY) mass is minimal in the PT (∼1mg) yet substantial and variable in the PH (∼20-40mg), presumably due to different uncinate process proportions in our tissue sections^49,50^ across all groups (***Figs.2B/C & S2K***). These findings readily recapitulate and refine the “histopathological dynamics” of T1D progression (including loss of ∼70-80% beta cell mass in T1DS^58^); they emphasize a profound alteration of the endocrine pancreas in the T1DL stage beyond the loss of beta cells; and they provide a general framework for our subsequent analyses focusing on ∼25,000 individual islets captured across the four donor cohorts.

### Islet endocrine hormone contents throughout T1D progression

Extending our quantification of pancreatic endocrine contents to individual islets reveals commonalities and differences compared to cumulative staining area assessments. A profound reduction of ProINS, INS and IAPP contents in T1DS islets is similarly presaged by a trend towards decreased ProINS and IAPP but not INS expression in AAb donors; T1DS and T1DL islets present with a “compensatory increase” of ProGCG, GCG and SST expression areas; and the PPY^+^ proportion of PH islets is variable and appears particularly large in T1DL donors (***Figs.2D & S2L***). At the same time, relative endocrine and CHGA areas remain constant throughout T1D progression thus corroborating, in conjunction with the ∼2.7-fold reduction of total CHGA staining area (***S2I***), a net islet loss in T1DL; and overall differences between donor groups tend to be more pronounced in PT than PH (***Figs.2D & S2L***). A direct comparison of PT and PH islets demonstrates that in all donor groups, despite comparable islet endocrine areas, PH islets express less CHGA, ProINS, INS, ProGCG, and GCG but equivalent IAPP and SST (***Fig.S3A***). While reduced alpha and beta cell mass in the PH of Ctrl donors has been noted earlier^50,59^, quantifying average islet hormone contents cannot resolve the potential contribution of different islet subsets to what appears an overall distinctive PH endocrine composition.

### Frequencies of islet subsets based on individual and combinatorial endocrine hormone expression

To evaluate how islet heterogeneity shapes net endocrine cell mass, we quantified islet frequencies based on their hormone content or lack thereof (defined as <1% hormone staining area of islet area). The significant increase of IDIs; ProINS^-^ or INS^-^) with disease development (T1DS PT: ∼64%, PH: 84%; T1DL PT/PH: >99%) constitutes the histopathological hallmark of T1D (***Figs.2E & S3B***). An unexpected feature of the non-diabetic pancreas revealed by recent 3D mapping is the notable abundance of small “GCG-deficient” islets^32^. We confirm this observation in our two-dimensional interrogation by demonstrating that ∼34% (PT) to ∼56% (PH) of Ctrl islets are “GCG-deficient”, and that ProINS^+^GCG^-^ islets are significantly smaller than ProINS^+^GCG^+^ islets (***Figs.2F/G & S3C/D***). The absence of alpha cells, as recently shown with human pseudo-islets exclusively composed of beta cells, does not appear to impinge on INS secretion dynamics^60^, suggesting that “GCG-deficient” islets constitute an integral part of endocrine pancreas physiology. Importantly, the ProINS^+^GCG^-^ islet fraction declines with T1D progression (including a significant reduction in AAb *vs.* Ctrl donor PTs), suggesting that this subset is particularly vulnerable in early T1D pathogenesis (***Figs.2G & S3D***).

Furthermore, a differential abundance of PPY^+^ islets in PT *vs.* PH aligns with cumulative PPY staining patterns (***Fig.2H***), and a subset stratification according to ProINS/PPY expression demonstrates that the Ctrl PH contains a ∼10% fraction of small PPY^+^ islets lacking beta cells that increases to ∼60% in T1DL (***Fig.2I***). Delta cells are found in ∼65% of islets, and their relative abundance is somewhat elevated in clinical T1D as noted before^61^ (***Figs.2J & S3E***). Lastly, we traced the evolution of hormone co-expression patterns. For alpha cells, a modest increase of Jaccard indices for CHGA, ProGCG and GCG with T1D progression is consistent with the relative rise of ProGCG/GCG content in islets afflicted by beta cell loss. For beta cells, Jaccard indices for relevant hormone combinations slightly decline in AAb donors before markedly plunging with T1D onset reflecting the profound perturbations of INS synthesis^62^ (***Fig.S3F***).

**Figure 2.**
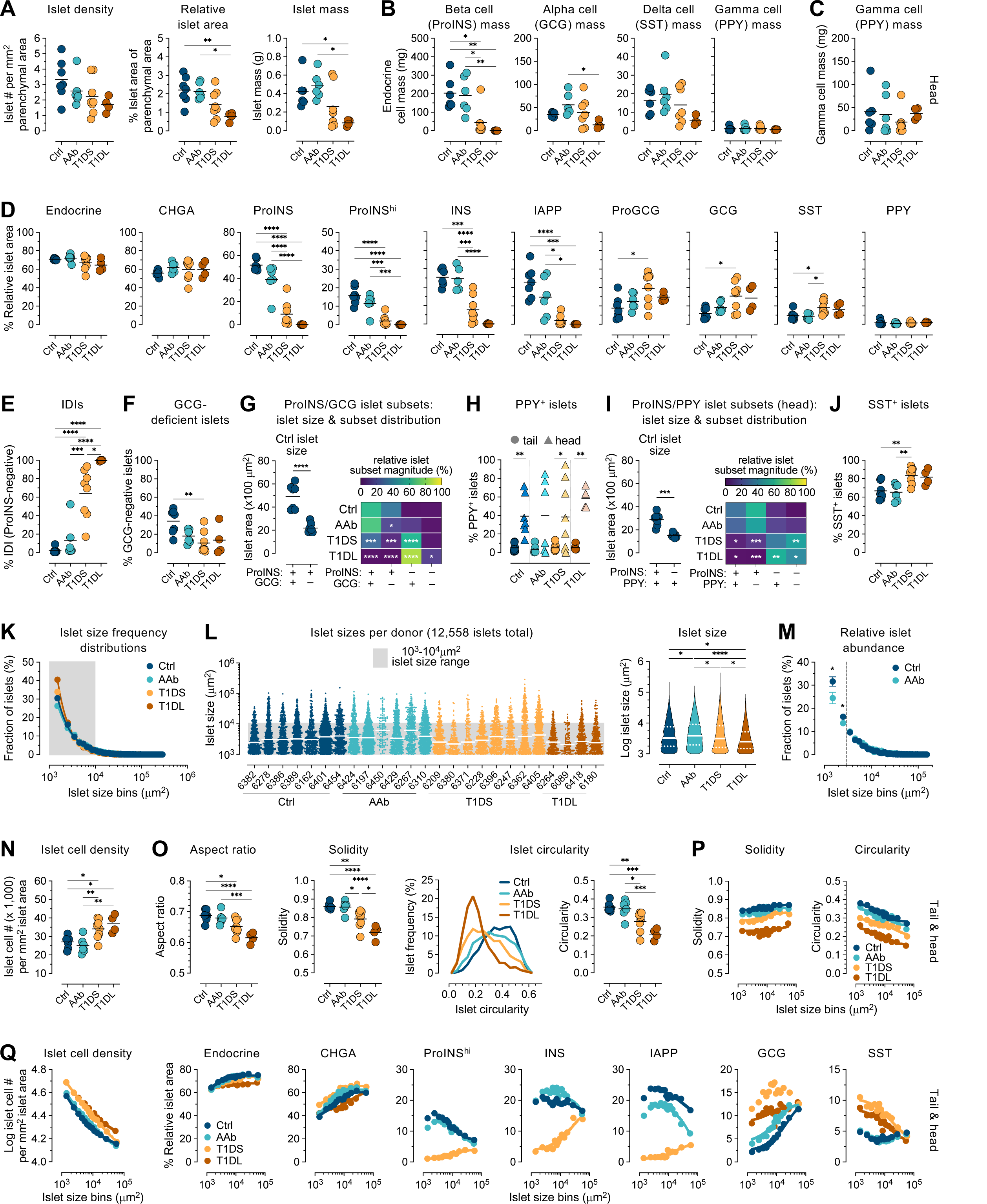
Pancreatic tissue section, islet and islet subset properties across stages of T1D progression. Unless noted otherwise, summary plots pertain to PT and display individual donor means (colored circles) and group means (black horizontal bars). **A.,** islet densities, relative areas and mass. **B./C.,** endocrine cell type mass in PT/PH was calculated from data in ***Fig.S2I*** and pancreas region weights (***Table S1***) excluding the two 5-year-old donors Ctrl 6382 and T1DS 6209. **D.,** fraction of indicated endocrine hormone staining areas in PT islets (“endocrine”: union of all hormone staining areas). **E.,** frequencies of IDIs (<1% ProINS content). **F.,** frequencies of GCG-deficient islets (<1% GCG content). **G.,** left: size of indicated islet subsets (individual donor medians and collective means); right: heatmap displaying relative magnitude of indicated ProINS/GCG islet subsets in each T1D stage (asterisks pertain to relative abundance of each islet subset in comparison to Ctrl donors). **H.,** frequencies of islets with ≥1% PPY content. **I.,** data display for combinatorial ProINS/PPY expression by PH islets as in panel G. **J.,** frequencies of islets with ≥1% SST content. **K.,** frequency distribution of islet sizes (colored circles: bin averages for all donors/group; all linear log-log curve fits: R^2^ >0.99). **L.,** size distribution of all 12,558 PT islets stratified according to donor group with individual donors ordered according to increasing age (white bars: medians); violin plot: islet size distributions (median/quartiles indicated) with statistical differences calculated using a mixed model. **M.**, relative abundance of bin-stratified islets from individual Ctrl *vs.* AAb donors (*c.f.,* panel K; vertical dashed line set at 3,000μm^2^; data are mean±SE). **N.,** geometric mean of islet cell densities. **O.**, islet aspect ratio, solidity and circularity; the binned histograms display donor group-specific islet circularity distributions. **P/Q.,** properties of PT/PH islets combined from all donors in indicated groups are displayed as a function of islet sizes stratified into 14 bins; additional parameters and goodness of exponential curve fits are detailed in ***Fig.S4A-H***. Due to notably weak CHGA and/or INS staining of PT but not PH sections from two donors (CHGA: Ctrl 6162; INS: Ctrl 6162, AAb 6450), respective data in panel D are excluded here. Statistical analyses were conducted with ordinary one-way ANOVA and Tukey’s multiple comparisons test (*p<0.05, **p<0.01, ***p<0.001, ****p<0.0001).

### Islet architecture and its histopathological alterations

The geometric properties of islets have mostly been studied in non-diabetic pancreata^63–66^ and in accordance with these reports, Ctrl islet areas span across two orders of magnitude (∼10^3^-10^5^μm^2^) following a power distribution that is notably skewed toward smaller islets (similar to reconstructed 3D analyses^64^, ∼80% of islets are found in the 10^3^-10^4^μm^2^ range) and roughly maintained throughout T1D progression (***Figs.2K/L & S3G/H***). Upon closer inspection, however, disease stage-specific deviations are discernible especially in the PT where islet size positively correlates with age (***Fig.S3I***). Here, mixed model analyses demonstrate a significantly increased median islet size in AAb *vs.* Ctrl donors that appears to result from the specific loss of islets <3,000μm^2^ (***Fig.2L/M***). Since such small islets constitute ∼48% of all islets in Ctrl subjects, a corresponding ∼1.6-fold islet density decrease in AAb donors, though not significant here, is notable nonetheless. A subsequent decline of median islet size in T1DS and especially T1DL donors (***Fig.2L***) is further accompanied by an increase of cellular densities in islets, confirming an earlier report^19^ and conceivably reflecting a local deprivation of trophic INS effects previously invoked as cause for exocrine pancreas shrinkage^57^ (***Fig.2N***). Although some of these patterns are not apparent in the PH, higher cellular densities in T1DS/T1DL islets are comparable for both PT/PH compartments (***Fig.S3G/H/J***). Thus, a T1D-associated decrease of islet size accompanies the reduction of overall islet densities (***Fig.1A***; see also ref.^67^) and compounds the loss of islet mass.

A recent 3D mapping of INS- and GCG-staining volumes in non-diabetic donors demonstrated that average human islet size is smaller than previously thought^32^. Working with similar assumptions about circularity and sphericity of islet objects, we used our islet area measurements to calculate median/mean islet diameters and volumes, collectively demonstrating that PT islets tend to be larger than PH islets (especially in AAb/T1DS but not T1DL donors) (***Fig.S3K***). Remarkably, the volume-based demonstration that islets have an average diameter of ∼65μm^32^ closely matches our estimates that median islet diameters in Ctrl donors range from ∼61μm (PH) to ∼64μm (PT). Similarly, while islets with >91μm diameter constituted ∼26% of all islets yet contributed ∼75% to beta cell mass in that report^32^, we find that islets above average size (>89μm diameter) account for ∼28% of islets (a proportion that is near identical across PT and PH as well as disease stages) and contribute ∼70% to islet mass in all donor groups (***Fig.S3L***).

In contrast to the subtle alterations of islet areas and a trend towards increased islet perimeters despite declining islet sizes with advancing disease (***Fig.S3M***), the progression of two-dimensional shape descriptors is profound: islet aspect ratio, solidity, and circularity significantly decline from Ctrl and AAb to T1DS and T1DL stage, a pattern that again is somewhat more accentuated for PT than PH islets (***Figs.2O & S3N***; aspect ratio, solidity and circularity are dimensionless measurements in the range of 0-1 that respectively describe object elongation, overall concavity, and similarity to a circular object; equations in ***Fig.1F***). Together, these alterations signify a stark deterioration of islet architecture. Our results indicate that inclusive whole-slide image analyses capture basic islet architectural and hormone expression features that readily align even with contemporary volumetric measurements^32,33^ and thus imbue the scope of observations reported here with greater confidence.

### A basic exponential relationship of islet size and islet properties

The heterogeneity of islet composition has long been recognized^58,68^ and in an attempt to account for disparate islet properties (including the conspicuous lack of alpha and/or beta cells specifically in smaller isles), we considered islet size as a practical organizing principle^69,70^. Correlating islet size frequency distributions in Ctrl pancreata with other islet features, we find that simple exponential relationships consistently capture these associations: increasing islet size correlates with greater islet cell numbers, solidity, endocrine/CHGA content and alpha cells but lower islet cell density, circularity as well as beta and gamma cell contents; these patterns only diverge for the smallest islets (<2,000μm^2^) with comparatively reduced beta cell fractions, as well as for SST content which is slightly elevated in both smaller and larger islets (***Figs.2P/Q & S4A/B***).

Several of these exponential relationships exhibit a striking resiliency in the face of T1D progression (endocrine and CHGA content) or are subject to a modest size-independent depression (solidity and circularity) or elevation (cellular density) indicative of pathogenetic processes encompassing islets of all sizes. While correlations of islet size with specific hormone abundance remain broadly similar in AAb *vs.* Ctrl donors, the T1DS stage features three deviations: a prominent inversion (overall reduced yet relatively higher ProINS, INS and IAPP expression by larger islets), a deterioration (ProGCG/GCG), and an emerging exponential association (SST decline as a function of islet size). In T1DL subjects, these patterns, apart from the profound beta cell loss, are mostly maintained or exacerbated (***Figs.2P/Q & S4C-H***). The dynamic regulation of these exponential relationships reflects an early loss of beta cells especially in smaller islets, preservation of residual beta cell mass in larger T1DS islets as well as the broad disappearance of beta cells and markedly altered islet composition in T1DL donors (***Fig.S4I***). Collectively, our semi-automated whole-slide image analyses confirm central tenets of T1D pathology and readily identify multiple previously un- or under-appreciated islet properties in normal pancreata and over the course of T1D.

### Integrated histopathology throughout T1D development and progression

The true analytical potential of our approach to characterization of islet heterogeneity lies in elucidating complex combinatorial property patterns. We therefore adapted a dimensionality reduction tool commonly employed for transcriptomic single-cell analyses and performed UMAP clustering^71^ of all ∼25,000 individual islets (rather than single-cells) captured in our four donor cohorts. This strategy generates five major clusters (I–V), four of which contain 3-4 subclusters (A-D), and a regional stratification illustrates that clusters III and IV are predominantly found in the PH, underscoring principal differences in the organization of endocrine compartments in PH and PT (***Fig.3A***).

**Figure 3.**
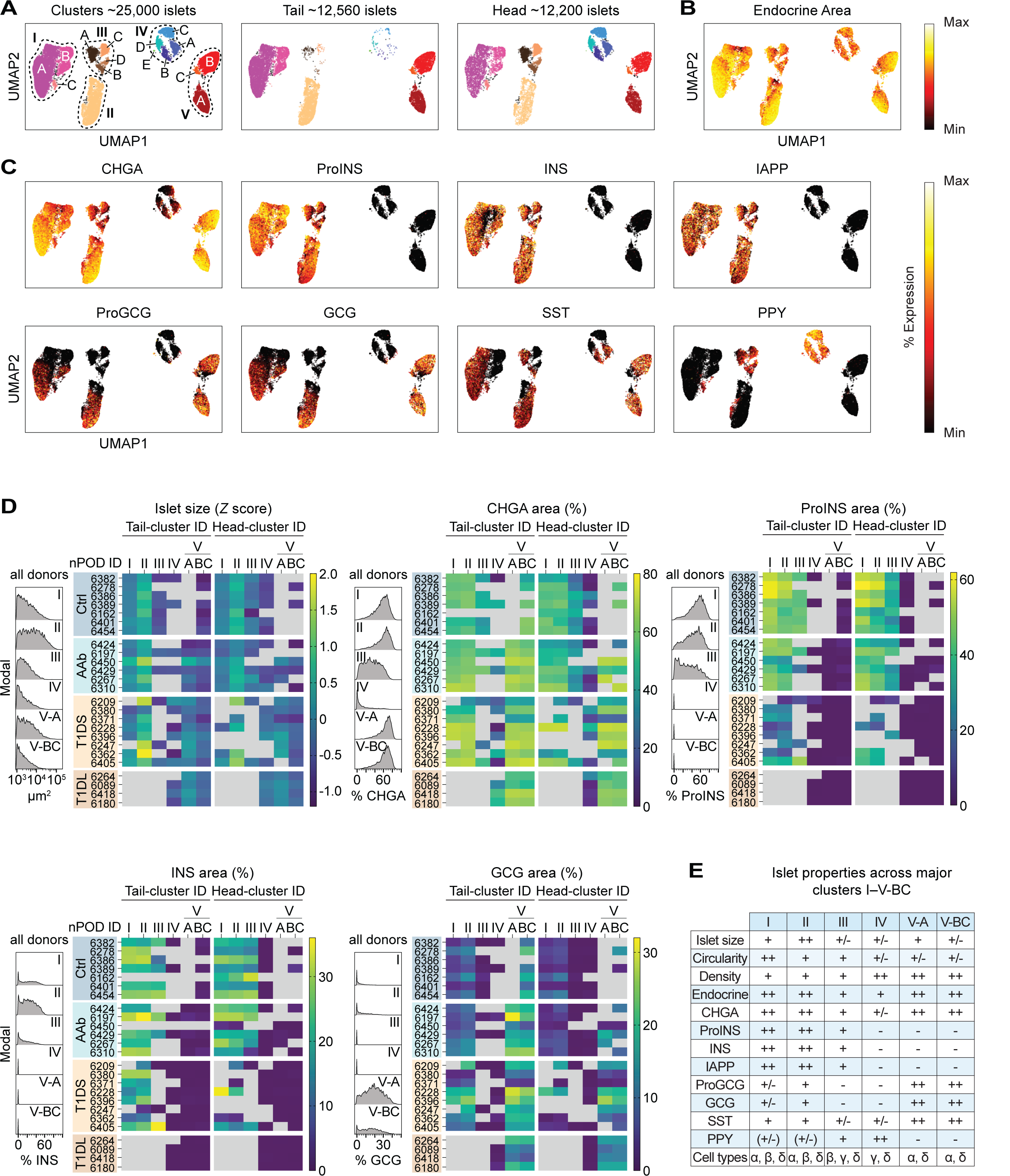
UMAP clustering of “single-islets”. **A.**, “single-islet” UMAP cluster annotation comprising all donor islets and stratification according to PT/PH regions (subcluster IV-E is sample-biased and excluded from further consideration). **B. & C.,** UMAP clustering of all islets with relative staining area percentages for individual endocrine hormones displayed as color gradient (min-max area: endocrine and CHGA: 0–100%; ProINS: 0-90%; INS: 0-50%; IAPP: 0-40%; ProGCG: 0-60%; GCG: 0-30%; SST: 0-20%; PPY: 0-100%). **D.,** histograms pertain to indicated UMAP clusters and display islet feature distributions combined from all donor PT/PH sections; the adjacent heatmaps stratify islet size and relative hormone staining areas (CHGA, ProINS, INS, GCG) across T1D stage, individual donors (listed in order of increasing age within each group), islet cluster affiliation, and PT/PH regions. Note that not all donors have islets populating each cluster and we further omitted values if clusters contained <3 islets or <2 donors; as in Fig.2, selected CHGA (Ctrl 6162) and INS (Ctrl 6162, AAb 6450) data were also excluded (missing/excluded values rendered in gray). **E.,** summary of distinctive islet properties across UMAP clusters.

Visualization of individual hormone expression levels across these clusters emphasizes both shared and distinct properties of cluster-constituent islets: endocrine areas are comparable for most clusters though slightly diminished for cluster IV, a pattern that is more pronounced for CHGA expression which is strongly reduced in cluster IV and somewhat decreased in cluster III (***Fig.3B/C***). Beta cell-containing islets are found exclusively in clusters I/II/III; alpha cells populate the majority but not all cluster I/II islets, are absent from cluster III and very infrequent in cluster IV, but a prominent presence in clusters V-A/V-BC; delta cells are distributed across all major clusters; and PPY^+^ islets, outside a smattering found in clusters I/II, reside in cluster III and dominate cluster IV (***Fig.3C***). The islet histograms and heatmaps in ***Figs.3D & S4J*** provide a more granular perspective on distinct cluster properties by resolving architectural and hormone expression properties at the level of individual donors, and are complemented by various scatter plot arrangements to illustrate more subtle differences and allow for the clarification of significant differences (***Fig.S5A-H***).

Cluster stratification shows a hierarchy of average islet sizes (cluster II>>I>V-A>III∼IV>V-BC) with near identical islet dimensions for respective PT and PH clusters (exception: AAb clusters I/II/V-BC where PT islets are bigger than PH islets) (***Figs.3D, S5A/C & S6A***) indicating that the trend towards larger islets in the PT must be grounded in regionally different cluster abundances. To wit, the PH of Ctrl donors contains a prominent contingent of small cluster IV islets (median diameter: 44μm) but combined cluster I-III insulin-containing islets (ICIs) have the same median size as PT islets (64μm diameter). A similar ranking of islet circularities (cluster I>II∼III>IV/V-A/V-BC) suggests that cluster I islets, due to their higher circularity and general preponderance in the non-diabetic pancreas (see below), may well have contributed to the historic notion of a “standard islet”^72^, and few differences in circularity or endocrine properties are observed for PT and PH islets within the same clusters (***Figs.3D, S5A/C & S6B***). Also, note the differential ProINS and IAPP content for clustered ICIs (I>II>III); absence of alpha/beta cells and low CHGA but high PPY expression in cluster IV islets; and elevated ProGCG/GCG expression levels of cluster V islets in conjunction with slightly increased SST content (***Figs.3D, S4J & S5B/D***). In agreement with the idea that INS may exert paracrine trophic effects, islet cell densities in clusters I/II exhibit an inverse correlation with beta cell content across disease progression while IDIs in PH cluster IV and the T1D-associated clusters V-A/BC are ∼35% denser than cluster I/II/III ICIs (***Figs.3D & S6C***). A succinct overview of these distinctive cluster properties is featured in ***Fig.3E***.

While UMAP analyses, by design, foreground discriminating cluster features, a comparison of islet properties within individual clusters across T1D stages also discloses some revealing differences: first, in transition from Ctrl to T1DS stage, ProINS and IAPP but less so INS content is progressively reduced in cluster I, II and III ICIs suggesting that modulation of ProINS and IAPP expression reports T1D pathogenetic processes with particular sensitivity (***Figs.S5F/H***). Second, opposing trajectories for islet size increase *vs.* circularity decrease in cluster II ICIs and especially cluster IV (PH) and V-A/V-BC IDIs can serve as a histopathological correlate for the dynamics of disease progression (***Figs.S5E/G***). Considering that larger Ctrl and AAb islets feature lesser ProINS/IAPP expression (***Figs.2Q & S4B/D***), our observations are consistent with islet size-dependent disease kinetics (*i.e.* early targeting of smaller ICIs and later appearance of larger IDIs).

### Islet cluster size redistribution as an early histopathological hallmark of T1D development

Arguably most relevant is the profound redistribution of relative cluster magnitudes with disease progression (***Fig.4A***). Heatmap (***Fig.4B***) and scatter plot (***Fig.4C***) displays demonstrate that cluster I/II islets account for the majority of islets in Ctrl pancreata (PT: cluster I ∼70%, II ∼25%; PH: cluster I ∼48%, II ∼17%; also III ∼25%; IV ∼10%). In AAb donors, this distribution is markedly altered due to a 39-45% reduction of cluster I islets accompanied by a compensatory 45-87% increase of cluster II islets (***Fig.4D***); this conclusion also applies to the individual collapse of subclusters I-A/B that are primarily distinguished by the presence/absence of delta cells (***Fig.S6D***), and cluster III islets in the PH including both SST^+^ and SST^-^ subsets (*i.e.,* cluster III-A/C reduction and cluster III-B/D increase) (***Fig.4E/F***) thus denoting the same fate for ICIs regardless of SST content.

**Figure 4.**
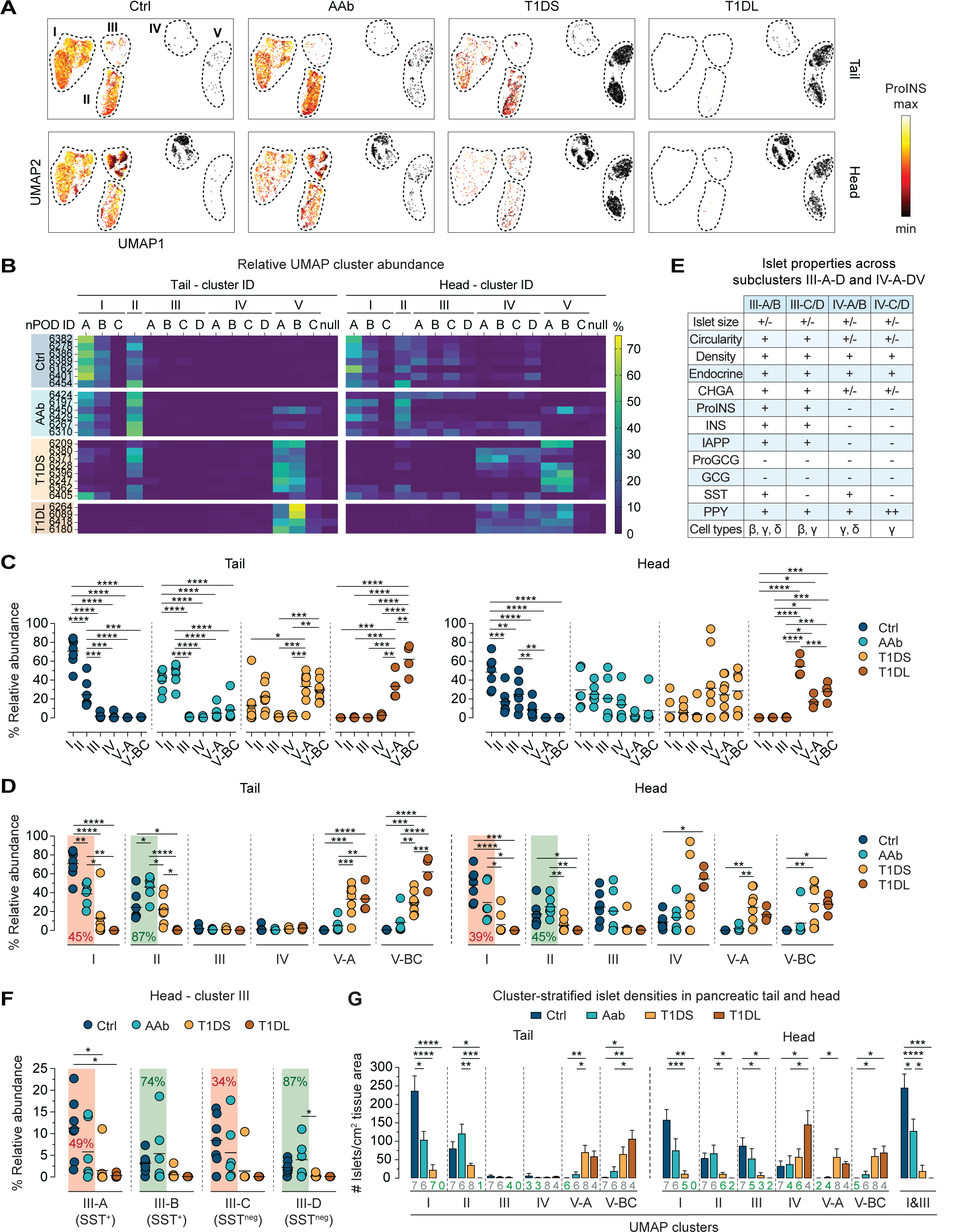
Redistribution of relative UMAP cluster sizes as a distinctive feature of early T1D development. **A.**, UMAP cluster display of all islets as a function of pancreas region and disease stage; the color gradient pertains to relative ProINS expression levels (min-max area: 0-90%). **B.,** heatmaps stratifying relative islet cluster sizes across T1D stages (individual donors listed in order of increasing age within each donor group), cluster affiliation (null, unclustered), and PT/PH regions. **C.,** relative cluster sizes in PT/PH within each donor group. **D.,** same data as in panel C now arranged for comparison of respective cluster magnitudes across T1D stages; Ctrl *vs.* AAb donors: note the relative decline of cluster I size means (red background) and accompanying increase of cluster II size means (green background). **E.,** summary of distinctive islet properties in PH subclusters III-A-D/IV-A-D (a further distinction of the four subcluster pairs according to differential immune cell burden is detailed in Fig.5F). **F.,** relative magnitudes of subclusters III-A-D in PH across T1D disease stages. **G.,** densities of cluster-stratified islets in parenchyma of PT (left) and PH (right; the far-right plot features combined cluster I/III islets since “redistribution dynamics” for these clusters are comparable, *cf.* panels D/F). Values below bars indicate the number of respective donors represented by the corresponding bar; in cases were not all donors are represented, the values are highlighted in green (data are mean±SE); for ANOVA details see Methods.

In transition to the AAb stage, cluster V islets emerge in some donors, and a progressive deterioration of cluster I/II magnitude in T1DS donors is accompanied by a robust increase of cluster V islets. In the PH, this pattern coincides with a relative growth of cluster IV (which “constitutively” lacks alpha/beta cells), including all of its SST^+^ and SST^-^ subclusters (***Figs.4D & S6E***). The reconfiguration of cluster sizes culminates in T1DL cases where IDIs in cluster V (PT) or clusters IV/V (PH) practically eclipse a minute fraction of residual cluster II/III ICIs (***Fig.4A-F***). Throughout T1D progression, distinctive differences between PT and PH cluster abundance (clusters I/II/V: PT>PH; clusters III/IV: PH>PT) remain broadly intact (***Fig.S6F***), indicating that despite regional differences, T1D pathogenesis proceeds in a somewhat synchronized fashion throughout the entire pancreas.

Translating the “dynamic regulation” of relative islet cluster magnitudes into the context of whole tissue sections illustrates the gradual progression of cluster-stratified islet densities with T1D stages, especially the significant reduction of AAb cluster I islet densities in the PT or of combined cluster I/III islet densities in the PH (***Fig.4G***). Furthermore, the conspicuous growth of PH cluster IV with disease progression is readily explained by cluster III ICIs that, after loss of beta cells, segregate together with cluster IV IDIs, thereby raising average islet size and lowering circularity (***Figs.4E/G & S5G***). Lastly, a donor-specific assessment of cluster frequency distributions (***Fig.4B***) permits the identification of several outliers and associated considerations, as detailed in the legend to ***Fig.S6F***.

### Insulitis and beyond: scope and focus of the immune cell burden

To quantify immune cell abundance, distribution, and islet association, we first counted CD45^+^ immune cells localized within (intra-islet) or immediately surrounding (peri-islet) individual islets (*cf., **Figs.1H & S1A***) to assess instances of insulitis (≥15 CD45 cells associated with ≥3 islets^73^). As shown in ***Fig.5A/B***, instances of insulitis (0/7 Ctrl, 3/6 AAb, 6/8 T1DS, 1/4 T1DL donors) are in overall concordance with prior manual determination by nPOD pathologists (***Table S1***) and constitute an expectedly rare occurrence^10^ in our T1DS cohort (1.3% [PH] to 3.2% [PT] of islets) that nevertheless is accompanied by a ∼2.5-fold increase of islet-associated CD45 cell numbers and densities. Though not significant, it is striking that a similar immune cell burden is already present at the AAb stage regardless of even lower insulitis frequencies (0.6% PH, 0.8% PT) (***Fig.5A/B***) and that its relative magnitude closely mirrors the average pancreatic increase of islet-reactive CD8^+^ T cells in AAb subjects^74^; considering specific CD8^+^ T cell frequencies in peripheral blood^74^, early immune cell recruitment to pancreatic islets may thus be mostly stochastic before enrichment of beta cell-specific CD8^+^ T cells becomes discernible at T1D onset. In the subsequent T1DL stage, islet infiltration has largely receded to levels comparable to Ctrl donors (***Fig.5A/B***).

**Figure 5.**
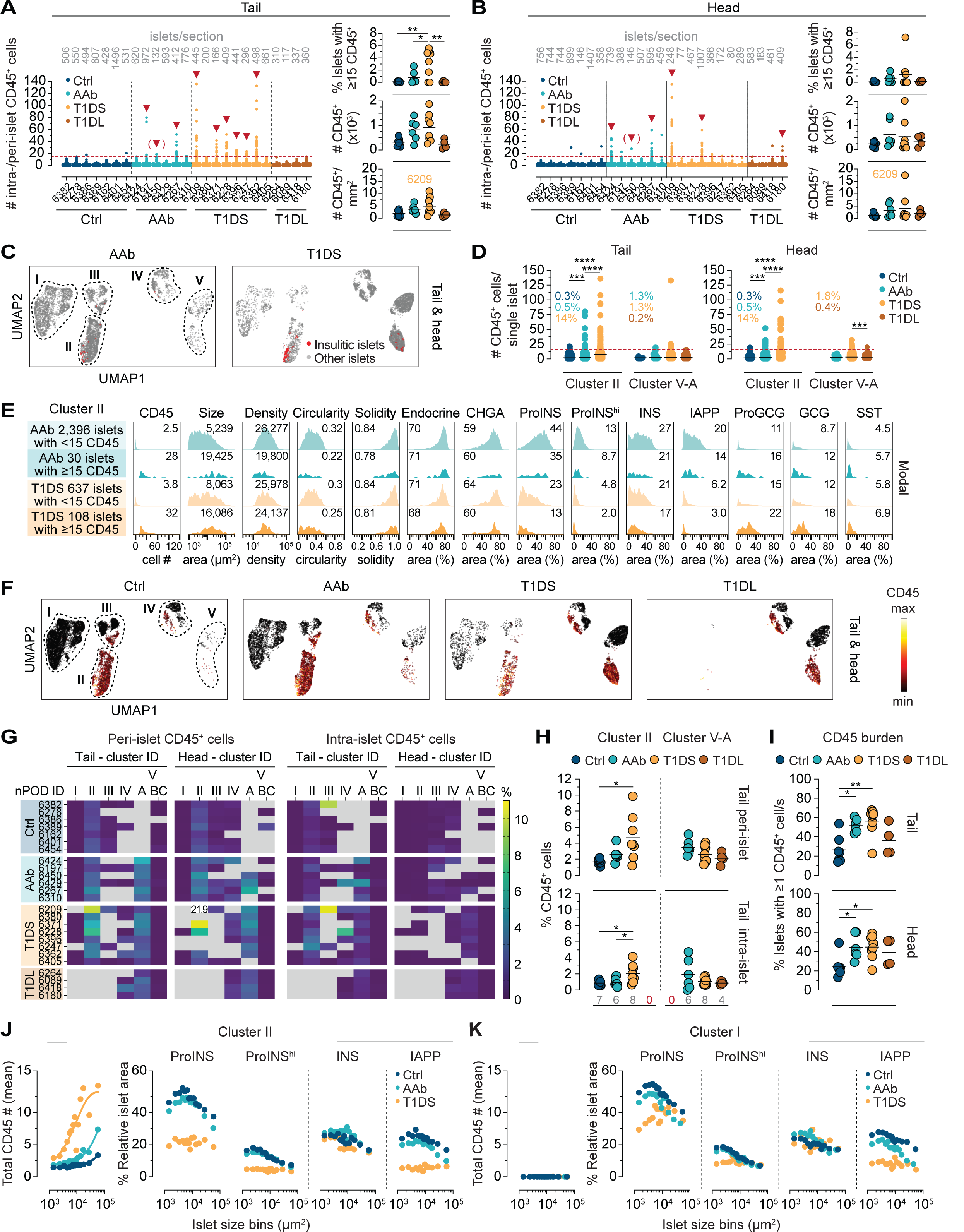
Insulitis and immune cell association with pancreatic islets. **A.**, left: total numbers of islet-associated CD45 cells (*i.e.,* intra- and peri-islet) for every single PT islet from all individual donors (listed in order of increasing age within each group; gray values on top are numbers of islets captured for each donor). The dashed red line indicates the “insulitis threshold” of ≥15 islet-associated CD45 cells, and red arrows highlight donors with ≥3 islets meeting the insulitis definition (AAb 6450: insulitis in two PT and one PH islet). Right: frequencies of insulitic islets (top), total number of islet-associated CD45 cells in tissue sections (middle), and islet-associated CD45 cell densities (CD45 numbers/mm^2^ peri-islet/islet area; bottom). **B.,** data display for PH islets as in panel A. **C.,** UMAPs depicting cluster localization of insulitic islets in AAb and T1DS donors. **D.,** absolute CD45 cell numbers associated with individual islets in clusters II ICIs and V-A IDIs of respective donor groups (each symbol represents an islet; dashed red line: “insulitis threshold”; color-coded values are percentages of insulitic islets in respective clusters of PT and PH). **E.,** properties of cluster II insulitic *vs.* other islets in AAb and T1DS donors (combined PT/PH islets from all AAb or T1DS donors; values are means, or medians for islet size/area; density: islet cells/mm^2^ islet area). **F.,** UMAPs of combined peri- and intra-islet CD45 cell frequencies (color gradient [min-max] CD45: 0-10%) across T1D stages in PT/PH. **G.,** heatmaps of average peri-/intra-islet CD45 cell frequencies stratified across donor group, individual donors listed in order of increasing age within each donor group, cluster affiliation, and PT/PH regions (note out-of-range value for T1DS 6209 cluster II peri-islet CD45 cell frequency; missing/excluded values in gray). **H.,** peri- and intra-islet CD45 cell frequencies in PT clusters II/V-A (values above x-axis are numbers of donors analyzed for each cluster and disease stage; T1DL and Ctrl donors lack cluster II and V-A islets, respectively). **I.,** frequencies of islets with ≥1 associated CD45 cell across disease stages in PT/PH. **J.,** combined PT/PH cluster II islets stratified across donor groups (Ctrl, AAb, T1DS) were “binned” according to islet size for display of associated CD45 burden and ProINS/INS/IAPP expression. **K.,** same as panel J but for cluster I islets. For ANOVA details in panels A, B, D, H and I, see Methods.

Our cluster stratification permits a refined assessment of insulitis distributions which are preferentially restricted to cluster II where 14% (PT) to 18% (PH) of islets are insulitic in T1DS cases. Such lesions are less pronounced in cluster V-A, very rare in clusters III/IV, and absent in clusters I/V-BC (***Fig.5C/D***). Although insulitis frequencies are lower in AAb donors, islet-associated CD45 counts in cluster II are higher than for Ctrl individuals comparing pooled donor groups, and an average of 2-3 CD45 cells per islet in cluster V-A of AAb, T1DS and T1DL donors constitutes a residual presence of immune cells no longer engaging beta cells (***Fig.5D***). Since cluster II islets histologically appear to constitute the principal target of autoimmune attack, we next compared the features of insulitic *vs.* non-insulitic islets in AAb and T1DS donors. Insulitic islets are noticeably larger, with a modest reduction in cellular density, circularity and solidity. Endocrine and CHGA contents remain unaltered while beta cell hormones are decreased, and alpha and delta cell hormone areas are comparatively increased (***Fig.5E***). Altogether, our findings demonstrate the utility of automated image analysis strategies for expedient insulitis diagnosis, elucidation of distinctive insulitic islet properties, and capture of the overall immune cell burden that is substantial in the AAb stage despite little insulitis.

### Association of immune cells with islets: from “physiological” to preclinical to pathological

In aggregate, *i.e.,* across all disease stages and pancreas regions, ∼52% of islet-associated immune cells are found in cluster II, ∼8% in cluster III, ∼13% in cluster IV, and ∼26% in cluster V-A. Notably, ∼40% of CD45 cells therefore associate with cluster IV and V-A islets despite their lack of beta cells. Since the use of absolute islet-associated CD45 cell counts in the consensus insulitis definition does not clearly account for islet size^73^, some investigators have adopted strategies for islet size correction such as quantifying immune cell numbers per mm^2^ islet area^23,26,75^. We employed a similar approach by quantifying islet-associated CD45 frequencies, namely the numbers of CD45 cells per nuclei in islets and peri-islet regions. Consistent with the overall cluster affiliation of islet-associated CD45 cells, islets with increased CD45 cell frequencies primarily localize to clusters II/V-A (in which every single islet is associated with ≥1 CD45 cell) and, to a lesser extent, also clusters III/IV (***Fig.5F/G***).

While these findings echo the distribution of insulitic islets, they further reveal a “physiological component” of immune cell infiltration since cluster II islets in Ctrl donors, accounting for ∼17% (PH) to ∼25% (PT) of all pancreatic islets, on average harbor ∼1.7% CD45 cells in the peri-islet region alone. This fraction increases by ∼3-fold in T1DS donors even under exclusion of T1DS 6209, a young child with notably high peri-islet infiltrates previously considered a more aggressive “endotype 1” T1D case^76,77^. Intra-islet CD45 frequencies tend to be lower but are also elevated 2.4-5.2-fold in the T1DS group, and similar considerations apply to PH cluster III islets (***Figs.5H & S6G***). In contrast, cluster IV/V-A IDIs follow a different pattern: higher peri-islet CD45 cell frequencies in AAb donors (PT: 3.5%, PH: 4.2%) decline to ∼2.1% in T1DL cases with less pronounced trends also observed for intra-islet CD45 cells (***Figs.5H & S6G***). In fact, considering that the ratios of peri-islet to intra-islet CD45 frequencies tend to rise in cluster II ICIs and fall in cluster V-A IDIs across T1D stages (***Fig.S6H***) may provide clues to better untangle the temporal progression of immune cell-mediated beta cell destruction. Finally, considerations of the entire spectrum of islet-associated immune cells permit the identification of an important histopathological correlate for stage 1/2 T1D: the frequency of islets associated with ≥1 CD45 cell constitutes a straightforward metric that readily distinguishes Ctrl (25%) from both AAb and T1DS (45-55%) donors altogether underscoring the significant immune cell recruitment to islets already in the AAb stage (***Fig.5I***).

### Immune cell:islet interactions and compromised islet composition in the histopathological absence of immune cells

A donor group-specific correlation between CD45 burden and islet properties in cluster II, the primary histopathological locus of CD45 infiltration, is confounded by the overall contingency of islet features, including CD45 cell numbers and frequencies, on islet size (***Figs.2P/Q, S4A-H & S6I***); instead, we compared “size-binned” cluster II islets across donor groups. Here, an increase of islet-associated CD45 numbers with disease progression appears to have little bearing on islet architecture or alpha and delta cell abundance (***Fig.S6J***). In contrast, ProINS and IAPP, but not INS, content is markedly reduced in T1DS *vs.* Ctrl and AAb donors, especially in smaller islets (***Fig.5J***). Even more striking is the observation that similar though attenuated patterns are also recorded for cluster I islets, which do not harbor any CD45 cells (***Figs.5K & S6J***). Cluster I islet impairment may arise following “exhaustion” of residual beta cells tasked with overwhelming metabolic demands^67,78^ or as a consequence of beta cell-targeted autoimmune attack^79^ not captured here due to its dynamic nature and/or the restricted plane of histological analysis; in all likelihood, both scenarios apply to varying degrees to different cluster I and other ICIs^6^.

Collectively, our cluster I/II islet evaluation indicates the lack of a straightforward correlation between CD45 cells and islet composition, suggesting that T1D pathogenesis is at once highly dynamic (*i.e.,* CD45 cells may “come and go” before complete beta cell destruction is achieved), initially targets islets below average size (compounding T1D symptomatology since smaller islets are capable of comparatively greater INS secretion^80,81^), and appears broadly synchronized since it affects the majority of islets (*i.e.,* ICIs present with reduced ProINS/IAPP expression even in the histological absence of immune cells). The utility of the insulitis concept and its proposed amendments notwithstanding^82^, considerations of the entire range of immune cell associations with pancreatic islets are therefore necessary to clarify progression of T1D autoimmune processes.

### Spatial distribution of islets throughout T1D pathogenesis

We employed three complementary approaches to assess the spatial distribution of islets in pancreatic tissue sections: modified Ripley’s K function, Delaunay triangulation, and direct visualization of cluster-associated islet localities. Ripley’s K function^83,84^ is a descriptive statistic for detecting deviations from spatial homogeneity, here the “aggregation” of islets in contrast to their random distribution patterns (***Fig.6A*** and Methods). In the PT of Ctrl donors, a modest ∼1.2-fold density enrichment is observed for islets across median radial distances of 600-1,500μm followed by a convergence toward random distributions for greater radii (***Fig.6B***). Thus, a relatively even distribution of islets throughout the PT readily aligns with recent observations obtained in 3D interrogations of non-diabetic pancreata^32^. While near identical islet distribution patterns are recorded for AAb donors, notable density enrichments occur in T1DS (∼1.8-fold, peaking at 1,000μm) and T1DL (∼2.6-fold, peaking at 600μm) with the right Ripley curve distribution tails still trending toward random distribution of islets, albeit less so at the T1DL stage. (***Figs.6B & S7A***). Similar islet distribution patterns over the disease course are also found in the PH. However, the unique anatomy of the uncinate process (see below) is responsible for more pronounced density enrichments in all donor groups (***Figs.6C & S7A***). Since islet locations are immutable, these “aggregations” reflect a progressive reduction of tissue regions with unperturbed islet densities that is driven by a relative loss of islets at shorter distances consistent with the overall decrease of islet densities, cumulative areas, and mass (***Figs.2A & S2H***). Lastly, these observations are confirmed by fractal dimension analyses^85^ as illustrated in ***Fig.S7B***.

**Figure 6.**
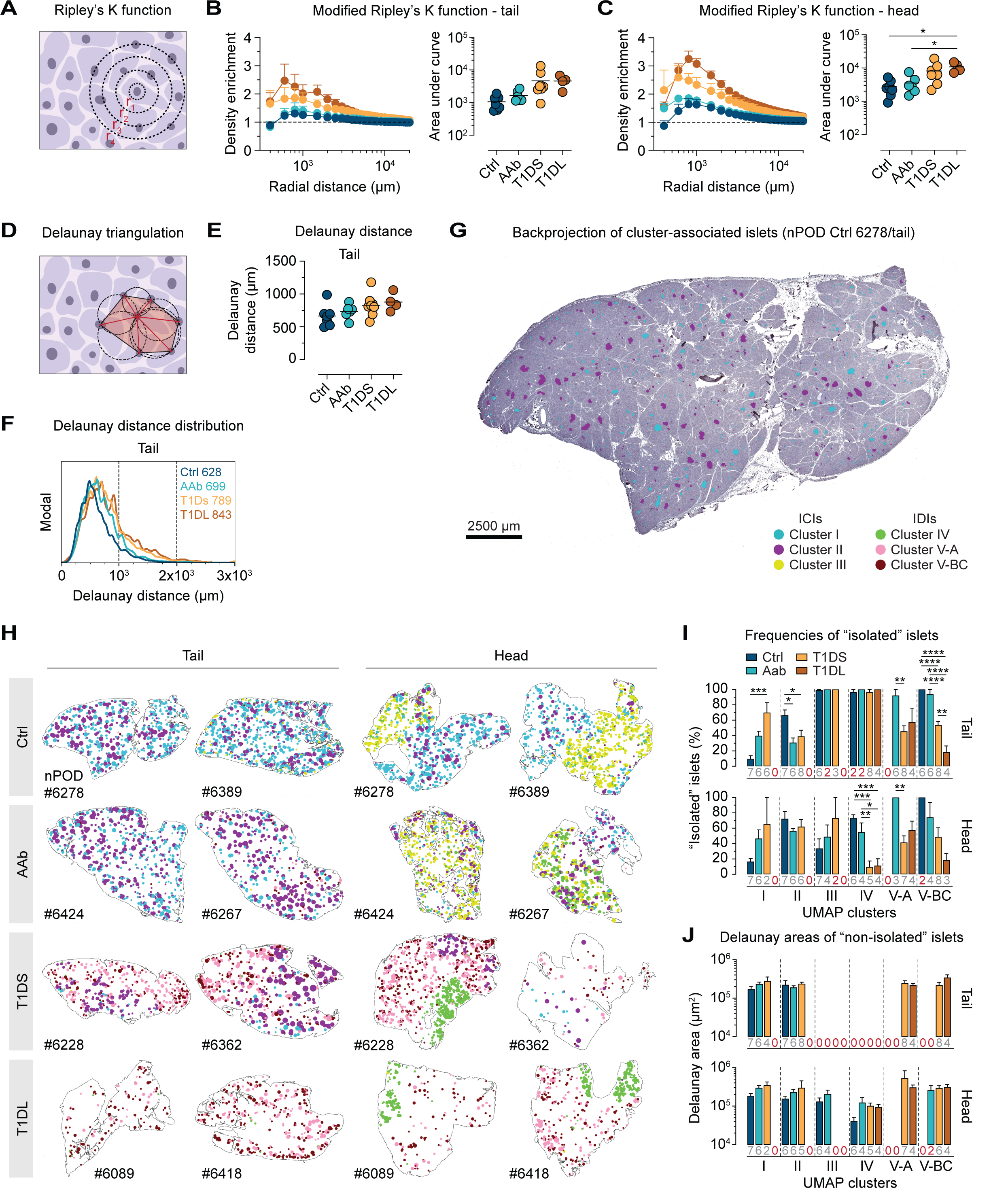
Spatial distribution of islets in the pancreas across the course of T1D pathogenesis. **A.**, Ripley’s K function: input values are radial distances between individual islets (center point) and expanding concentric circles that serve as bins to capture surrounding islets (other points) at increasing radial distances (see Methods for details). **B.,** left: islet density enrichments in PT of all groups as a function of radial islet distances ranging from 4×10^2^-2×10^4^μm; horizontal dashed line indicates random islet distribution (data are mean±SE). Right: areas under Ripley curve. **C.,** data display for PH islets as in panel B (T1DS 6380 excluded due to very scarce islets, *cf. **Fig.S8A***). **D.,** Delaunay triangulation of a set of points/islets and associated circumcircles (gray); the center point/islet is connected by six red triangle sides to other points/islets and the average length of these lines is the mean Delaunay distance; the mean area of the six shaded triangles is the mean Delaunay area. **E.,** mean Delaunay distances between islets across T1D progression in PT. **F.,** distribution of mean Delaunay distances according to T1D stage. **G.,** *in situ* projection of islets color-coded by cluster onto representative PT section. **H.,** pancreatic tissue section outlines populated with islets in their original location color-coded according to UMAP cluster affiliation; for facilitated visual presentation, islets are rendered circular and enlarged with relative size differences preserved (color legend in panel G). **I. & J.,** Delaunay triangulation was performed separately for islets in each cluster; islets with <2 neighboring islets within an 8mm distance are “isolated islets”. I., fraction of “isolated islets”. J., mean Delaunay areas of “non-isolated” islets. The values below each bar are the numbers of donors analyzed for each cluster and disease stage (“0”: absent values or data exclusion if <3 islets or <2 donors were represented in a given cluster; instances with 2 donors only are highlighted in red to indicate need for interpretative caution; data are mean±SE). For ANOVA details in panels B, C and E, see Methods.

### Progressive changes to islet neighborhoods

Delaunay triangulation organizes a set of points (islets) in a plane (tissue sections) into triangles whose circumcircles do not contain any of the points (and whose circumcenters constitute the vertices in Voronoi tessellation)^86^; thus, mean Delaunay distances and triangle areas can serve as a more local metric for average adjacency between individual islets and their proximate neighbors (***Fig.6D***). Here, Delaunay distances for PT islets increase by ∼30% from Ctrl to T1DL stage and display a similar though blunted trajectory in the PH (***Figs.6E* & *S7C***). At the same time, mean Delaunay distances are normally distributed in Ctrl pancreata with a slight extension of the right distribution tail representing islets that are more distant to each other; with T1D progression, the respective right distribution tails increase in particular in the PT (***Fig.6F* & *S7C***), and further alterations can be discerned by a cluster-level evaluation of Delaunay distances as detailed in ***Fig.S7D-G***. Collectively, our observations support the notion that the typically regionalized patterns of T1D histopathology^10^ represent residuals of a tissue-wide islet depletion, the extent of which is partially obscured by its fundamentally dispersed nature.

### Cluster-specific islet distribution patterns and the “dynamics” of T1D progression

To visualize the above relations more directly, we “projected” cluster-stratified islets back onto their original locations in the tissue sections (***Figs.6G & S7H***), revealing patterns unique to specific T1D stages: in the Ctrl PT, a relatively uniform distribution of cluster I islets is accompanied by a sparser presence of cluster II islets, a balance that in AAb donors shifts to visibly fewer cluster I islets. This pattern changes more decisively in T1DS cases where the appearance of predominantly cluster V-A islets crowds out remaining cluster II islet foci, and the process is completed in T1DL with the prominent emergence of cluster V-BC islets that often appear to populate the tissue area periphery (***Figs.6H & S8A***). In the PH, a similar progression pertains to the redistribution of cluster I/II/V islets, yet the most distinctive aspect is a striking regionalization of cluster III/IV islets representing the uncinate process (***Figs.6H & S8A***).

We employed “islet cluster-specific” Delaunay triangulations to quantify these impressions. First, “isolated islets” were defined as islets for which, due to fewer than two islets from the same cluster within an 8mm distance, Delaunay triangulation was not possible. As cluster I magnitudes collapse from Ctrl to T1DS stage (***Fig.4G***), the fraction of “isolated islets” increases (***Fig.6I***). At the same time, “isolated islet” frequencies in cluster II decline somewhat, reflecting a relative condensation of residual cluster islands that are in the process of disappearing with T1D onset (***Figs.4G & 6I***). The sparse cluster III/IV islets in the PT are practically all isolated and remain so throughout all disease stages. In contrast, the more abundant cluster III ICIs in the PH become increasingly isolated as their transition to IDI status following beta cell loss leads to consolidation with cluster IV islets (*c.f.,* ***Fig.4E/G***); in turn, the resultant growth of cluster IV is accompanied by a steep decline of “isolated islets” therein (***Fig.6I***). “Isolated islets” in clusters V-A/BC dominate at first appearance in the Ctrl and/or AAb stage, decline in T1DS with expansion of respective cluster magnitudes (***Fig.4G***), and for cluster V-BC are further reduced in T1DS (***Fig.6I***). The combined “dynamics” of these changes indicate that cluster V-A represents a transitional and cluster V-BC a terminal fate.

Second, quantification of Delaunay areas for remaining “non-isolated islets” reveals trends that partially mirror the “isolated islet dynamics” (*e.g.,* increasing Delaunay areas for cluster I islets), yet no significant differences emerge across disease stages for any of the clustered islet neighborhoods (***Fig.6J***). Altogether, these observations reinforce the notion that expanding areas of islet loss, increasingly populated by growing “diabetic” cluster V islands, encroach on progressively shrinking cluster I/II/III islands without a major disruption of the fundamental spatial organization of islet cluster regions.

### Insulitis and immune cell burden in spatial context

Finally, we leveraged the “back-projection” of islet subsets to visualize the neighborhood context of insulitic lesions. Using a color gradient to display the numbers of islet-associated immune cells, the resultant images demonstrate that rare instances of insulitis in AAb and T1DS donors often are the foci of larger neighborhoods with an overall increased CD45 cell burden; this pattern tends to be more pronounced for the PT than PH (***Figs.7A & S8B***). And lastly, our spatial data visualization also clarifies the distinctive pathology of three previously discussed “outlier” cases and is detailed in ***Fig.S8A/B***.

### Conclusions

This study combines multiplexed brightfield IHC of human pancreatic tissue sections, high-magnification whole-slide imaging, digital pathology, and development of a semi-automated image analysis pipeline implemented in the open-source pathology platform QuPath^37,38^ to trace the evolution of T1D development and progression. Integration of these modalities readily confirms the central tenets of pancreatic T1D histopathology and reveals multiple novel aspects about the dynamic organization of the pancreas in health and T1D disease, including the utility of islet size distribution as an organizing principle for islet heterogeneity, and an essentially identical endocrine organization across pancreas regions that locates residual differences specifically to the uncinate process in the PH. Notably, we define a series of new histopathological correlates for the stage 1/2 pancreas (*i.e.,* higher frequencies of islets with ≥1 associated immune cell; early targeting of small, including “GCG-deficient” islets; relative reduction of total ProINS/IAPP areas; pronounced collapse of islet cluster I magnitude) that position AAb subjects on the cusp of developing the very histopathological hallmarks that are distinctive for T1D; we demonstrate the absence of a direct correlation between islet composition and associated immune cell abundance (*e.g.,* decreased ProINS/IAPP content even in smaller T1DS ICIs without observable CD45 cells) thus emphasizing the importance to consider the entire spectrum of immune cell associations with islets beyond those affected by insulitis; and we foreground the “negative spaces” of T1D histopathology, namely the apparent loss of whole islets from expanding pancreas regions accompanied by a contraction of residual areas with “normal” islet density suggesting a multifocal and somewhat synchronized origin and progression of T1D pathology. We further submit that “single-islet” UMAP clustering can aid in better resolving histopathological properties and processes, and since our specific input variables may not be available for other studies, we provide a simple key that permits an approximate “clustering” of islets based on standard three-parameter stains (INS, GCG, CD45; PPY visualization may be added to distinguish cluster III/IV islets in the PH) (***Fig.S7I***). Altogether, we propose a revised natural history for T1D (***Fig.7B***) that may be leveraged for more targeted investigative tasks seeking to elucidate the autoimmune pathogenesis of T1D as well as the pathology-informed consideration of interventional modalities.

**Figure 7.**
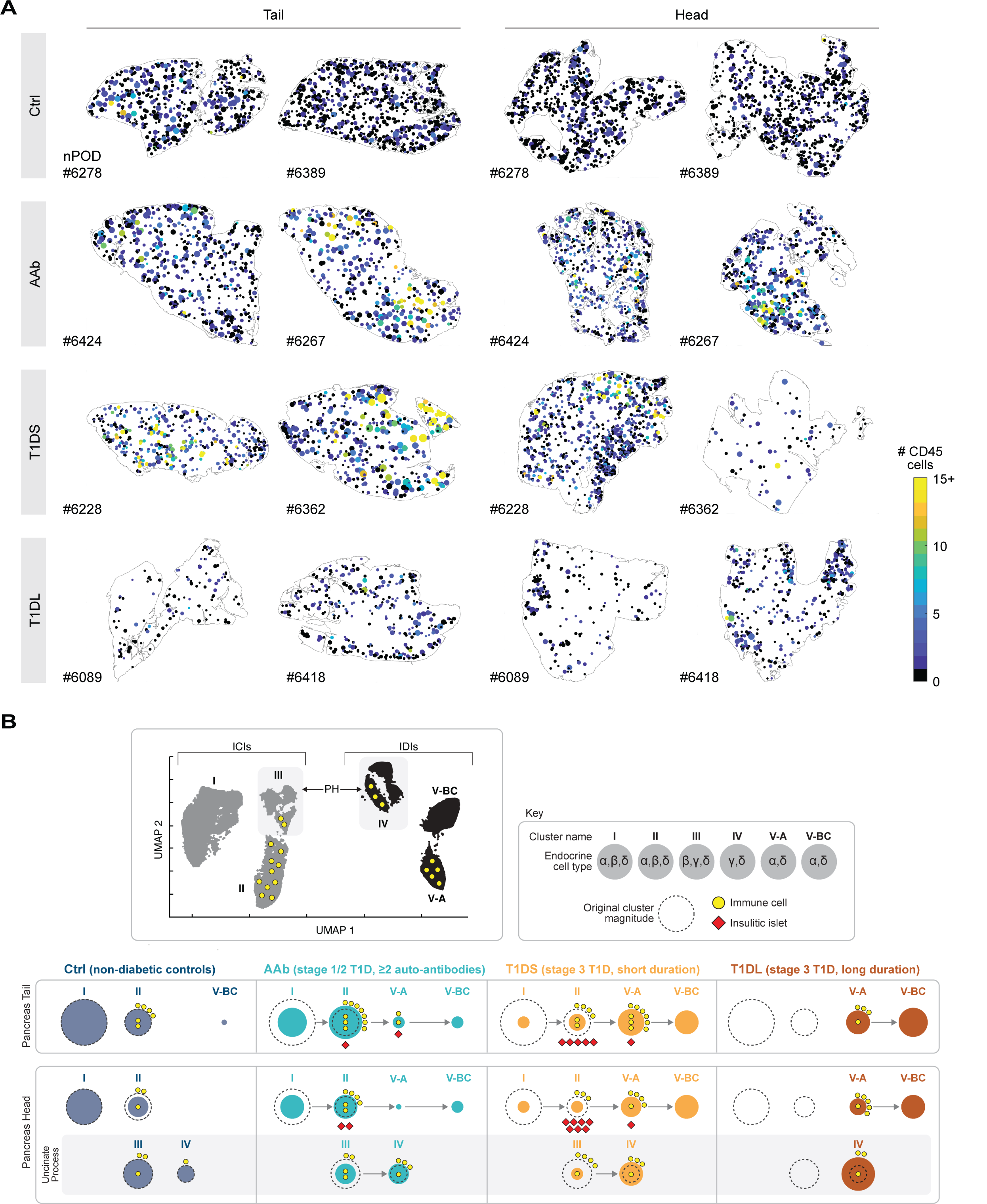
Spatial context for islet-associated immune cell burden & model summary. **A.**, pancreatic tissue section outlines (regions, T1D stage and nPOD donor IDs indicated) populated with islets color-coded according to the numbers (0-15+) of associated CD45 cells (islets are rendered circular and enlarged with relative size differences preserved). **B.,** summary model: islets are allocated to UMAP clusters I – V-BC according to geometric properties, endocrine cell content, association with immune cells, and Delaunay area. Top left: major cluster properties of ICIs (gray) and IDIs (black); top right: legend. Bottom: T1D progression in the PT and PH (divided into non-uncinate and uncinate process regions). The size of colored circles represents the relative magnitude of respective islet clusters across T1D stages; their changing sizes across disease progression is indicated by arrows between adjacent circles; the dashed perimeter lines indicate the original magnitude of clusters I, II, III and IV in Ctrl donors and serve as a visual reference for cluster size changes with disease progression (a further net reduction of islet cluster magnitudes due to loss of pancreas weight is not captured here; also note that cluster III islets, upon losing beta cells, become “re-classified” as cluster IV islets).

## LIMITATIONS OF THE STUDY

We note several limitations of the present study: the relative paucity of donor tissues compounded by donor variability, a perennial challenge for T1D histopathology studies; our analytical exclusion of small endocrine clusters and single cells (islet objects <1,000μm^2^) the fate of which is gaining increased attention^87,88^ and constitutes a focus of ongoing investigations; and the fact that refined analysis modalities and novel observations reported here, though grounded in effective capture of known histopathological hallmarks of T1D progression, will require validation in independent studies and/or larger cohorts. Our documentation of islet mass reduction in clinical T1D may be confounded by the possibility that individuals prone to T1D development have inherently smaller pancreata, a contention that at present, however, remains speculative^6^. Lastly, age-matching of pancreas donors is critical to our study design but implies different ages for disease onset (T1DL<T1DS<stage 1/2 T1D); additional histopathological investigations should therefore include donors matched for age-of-onset to better account for age-dependent variables in T1D pathogenesis.

## ACKNOWLEDGMENTS

We wish to thank Dr. R. Brody (ISMMS Biorepository and Pathology CoRE) for provision of additional pancreatic tissue sections used for MICSSS optimization, Dr. O. Madsen (Novo Nordisk) for the gift of ProINS antibody, Dr. P. Bankhead (University of Edinburgh) for advice about QuPath customization, Dr. E. Bagiella (ISMMS Center for Biostatistics) for assistance with statistical analyses, Dr. B. Rosenberg for advice about UMAP clustering, J. Gregory for design of the model figure panel, and Dr. S. Richardson for detailed feedback on the manuscript. This research was supported by Juvenile Diabetes Research Foundation Postdoctoral Fellowship JDRF 3-PDF-2018-575-A-N and a supplement (V.v.d.H.); Chan Zuckerberg Initiative grant DAF2019-198153 (S.M.); NIH grants CA224319, DK124165, CA234212, and CA196521 (S.G.); and NIH grants R01AI134971, R01DK130425, R21ES027916 and P30DK020541 (D.H.). To provide context for the multiplicity of observations reported here, we repeatedly cite consensus opinions, or the lack thereof, summarized in authoritative reviews; we apologize to the authors whose pertinent primary contributions are not explicitly mentioned here. Most importantly, we wish to thank the families of the organ donors for the gift of tissues.

## AUTHOR CONTRIBUTIONS

Conceptualization: V.v.d.H. and D.H.; investigation: V.v.d.H.; formal analysis: V.v.d.H., S.M., Z.M., K.C. and D.H.; software: S.M. and M.N.; writing - original draft: D.H.; writing - review and editing: all authors; visualization: V.v.d.H., S.M. and D.H.; resources: S.G., Z.M., A.L.P., I.K. and M.A.; funding acquisition: V.v.d.H. and D.H.; supervision: D.H.

## DECLARATION OF GENERATIVE AI AND AI-ASSISTED TECHNOLOGIES

No generative AI or AI-assisted tools or services were used at any stage of the writing process.

## SUPPLEMENTAL INFORMATION

**Supplementary Figures S1 – S8. *Figs.S1 – S8*** feature additional data related to ***Figs.1* – *7***.

**Supplementary Tables S1 – S3. *Table S1***, related to ***Table 1***, provides detailed information about pancreas specimens, demographic and clinical donor metadata, and HLA haplotype T1D risk; ***Table S2*** summarizes and stratifies all properties of the ∼25,000 individual islets captured in the present study; ***Table S3*** features information about antibodies and MICSSS staining conditions.

**Table 1.**
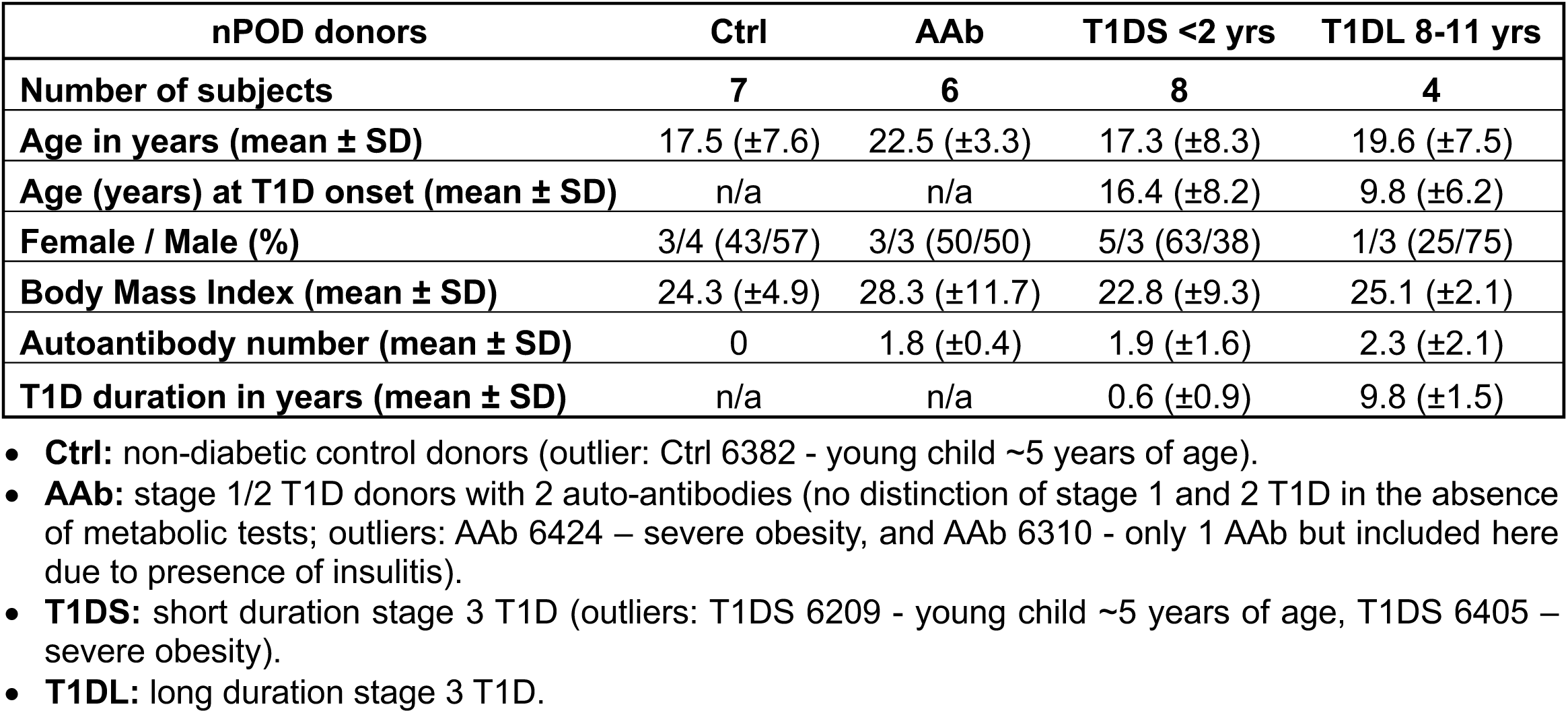
Overview of pancreatic organ donors. For further pancreatic specimen and organ donor details, see ***Table S1***.

| **nPOD donors** | **Ctrl** | **AAb** | **T1DS <2 yrs** | **T1DL 8-11 yrs** |
| --- | --- | --- | --- | --- |
| **Number of subjects** | **7** | **6** | **8** | **4** |
| **Age in years (mean ± SD)** | 17.5 (±7.6) | 22.5 (±3.3) | 17.3 (±8.3) | 19.6 (±7.5) |
| **Age (years) at T1D onset (mean ± SD)** | n/a | n/a | 16.4 (±8.2) | 9.8 (±6.2) |
| **Female / Male (%)** | 3/4 (43/57) | 3/3 (50/50) | 5/3 (63/38) | 1/3 (25/75) |
| **Body Mass Index (mean ± SD)** | 24.3 (±4.9) | 28.3 (±11.7) | 22.8 (±9.3) | 25.1 (±2.1) |
| **Autoantibody number (mean ± SD)** | 0 | 1.8 (±0.4) | 1.9 (±1.6) | 2.3 (±2.1) |
| **T1D duration in years (mean ± SD)** | n/a | n/a | 0.6 (±0.9) | 9.8 (±1.5) |

- **Ctrl:** non-diabetic control donors (outlier: Ctrl 6382 - young child ~5 years of age). - **AAb:** stage 1/2 T1D donors with 2 auto-antibodies (no distinction of stage 1 and 2 T1D in the absence of metabolic tests; outliers: AAb 6424 – severe obesity, and AAb 6310 - only 1 AAb but included here due to presence of insulinitis). - **T1DS:** short duration stage 3 T1D (outliers: T1DS 6209 - young child ~5 years of age, T1DS 6405 – severe obesity). - **T1DL:** long duration stage 3 T1D.

## SUPPLEMENTARY TABLES & FIGURE LEGENDS

**Supplementary Table S1. Pancreas specimen information & donor metadata.** All pancreatic tissue sections were provided by the Network for Pancreatic Organ Donors with Diabetes (nPOD). ***Table S1*** summarizes pancreas donor information arranged into four groups (Ctrl, AAb, T1DS, T1DL; within each group, donors are listed in order of increasing age) and comprises nPOD donor ID; formalin-fixed, paraffin-embedded (FFPE) tissue block details; donor demographics (gender, ethnicity, age, age at T1D onset, T1D duration); clinical parameters (body mass index, C-peptide, HbA1c, number and type of AAbs, cause of death); pathology assessments (pancreas weight, presence/absence of insulitis); and genetics (available HLA haplotypes were used in conjunction with ethnicity to calculate genetic T1D risk as detailed in Methods).

**Supplementary Table S2. Properties of individual islets donors, UMAP clusters across pancreas regions. *Table S2*** summarizes all properties of ∼25,000 individual islets captured in the present study and stratified according to donor group (including demographic and clinical metadata), UMAP cluster affiliation (I - V-BC) and pancreas region (PT, PH). ***Table S2*** *is not appended to the initial submission*.

**Supplementary Table S3. Antibodies and MICSSS staining conditions.**

**Supplementary Figure S1. MICSSS staining and semi-automated image analysis of pancreatic tissue sections. A.,** sequential MICSSS staining of eight pancreatic endocrine hormones and CD45. Each image pair features the native IHC image (left, number in upper left corner indicates position in staining sequence) and the same image overlaid with automatically captured staining area traces (right) and islet perimeter (white) (note that ProINS images are duplicated here from ***Fig.1G***, and islet #86 does not contain PPY^+^ gamma cells). Automated CD45 cell captures (highlighted in yellow) are restricted to areas within the peri-islet border (pink; set at a distance of 20μm from islet perimeter) since classifier training did not include immune cells located outside that border. **B.,** to assess the relative extent of staining overlap between INS or GCG and other endocrine hormones, we determined respective Jaccard indices by dividing the size of the staining area intersection by the size of the union of staining areas (expressed as fraction [%] of staining area) for each combination of hormone staining pairs; asterisks indicate statistical significance (*p<0.05, **p<0.01, ***p<0.001 and ****p<0.0001) in one sample t tests for values >2% staining overlap. **C.,** CHGA, INS, GCG and SST brightfield stains (brown) of PPY^+^ islet #206 (*c.f.,* ***Fig.1I***); note the absence of alpha and beta cells as well as little CHGA expression.

**Supplementary Figure S2. Pancreatic donor, tissue and islet properties across stages of T1D progression. A.,** HLA-II haplotypes were “binned” according to ref.^52^ and assigned the following numerical “risk values”: protective = −1, neutral = 0, moderate risk = +1, high risk = +2. **B. & C.,** correlation of donor age, BMI, pancreas weight and age of T1D onset as well as C-peptide and HbA1c. Goodness of linear or exponential curve fit (R^2^) is indicated, and asterisks indicate statistical significance of slope deviation from zero. “Outliers” as based on severe obesity (AAb 6424, T1DS 6405), very young age (Ctrl 6382, T1DS 6209) and comparatively better glucose handling (T1DS 6405) are indicated (HbA1c values for Ctrl 6162, AAb 6310, T1DS 6209, T1DS 6247 and T1DL 6180 were not available). **D.,** relative pancreatic weight (RPW: pancreas weight [g] / body weight [kg]) was calculated for the entire organ (left) as well as PT (middle) and PH (right) regions. **E. & F.,** absolute and relative parenchymal tissue section areas, and correlation of former with donor age in PT but not PH. **G. & H.,** absolute numbers of islets in PT and PH tissue sections as well as islet densities (numbers of islets per mm^2^ parenchymal tissue area), cumulative islet areas and islet mass in PH. **I.,** fraction of indicated total endocrine hormone staining areas normalized to parenchymal tissue areas in the PT (top) and PH (bottom). A reduction of ProINS areas in AAb donors would appear to contradict a reported elevation of total ProINS expression and ProINS:INS ratios in AAb *vs.* Ctrl donors^15^. However, those outcomes are in part dependent on the capture of small endocrine clusters (<1,000μm^2^) excluded in the present analyses; they appear to be more pronounced in single than double AAb^+^ donors^15^; and ProINS:INS expression ratios become exceedingly variable with disease advancement^15,89,90^. Thus, our observations do not challenge the emerging paradigm of altered beta cell prohormone processing in T1D progression^62^. **J.,** INS:GCG expression ratios were calculated with individual donor values featured in panel I. **K.,** endocrine cell type mass was calculated with respective PT and PH data in panel I. and corresponding PH weights (***Table S1***) under exclusion of the two 5-year-old donors Ctrl 6382 and T1DS 6209; Ctrl case 6405 is indicated due to their high alpha and beta cell mass. **L.,** fraction of indicated relative endocrine hormone staining areas in individual islets of the PH (“endocrine” refers to the union of all hormone staining areas). Due to notably weak CHGA and/or INS staining of PT but not PH sections from two donors (CHGA: Ctrl 6162; INS: Ctrl 6162, AAb 6450), the respective PT data are excluded in panels I.-K. Unless noted otherwise, data are scatter and mean with statistical analyses conducted using ordinary one-way ANOVA and Tukey’s multiple comparisons test adhering to the following convention: *p<0.05, **p<0.01, ***p<0.001, ****p<0.0001; ns, non-significant.

**Supplementary Figure S3. Properties of islets and islet subsets across stages of T1D progression. A.,** for comparative purposes, panel A. features the same data as in ***Figs.2D & S2L*** to contrast relative islet hormone staining areas in PT *vs*. PH. **B.,** frequencies of IDIs with <1% ProINS or <0.1% INS expression. **C.,** frequencies of islets with <1% GCG expression. **D.,** left: size of indicated PH islet subsets (individual donor medians and collective means); right: heatmap displaying the relative magnitude of indicated ProINS/GCG PH islet subsets in each T1D stage (asterisks pertain to relative abundance of each islet subset in comparison to Ctrl donors)**. E.,** frequencies of islets with ≥1% SST expression. **F.,** Jaccard indices were calculated as detailed in Methods for dual combinations of hormones co-expressed by alpha cells (CHGA, ProGCG, GCG) or beta cells (CHGA, ProINS, ProINS^hi^, INS, IAPP) and are displayed as a function of T1D stage. The reduction of ProINS-INS Jaccard indices with disease progression appears to contravene the reported increase of ProINS/INS co-localization in T1DS donors. However, the latter phenomenon was largely restricted to young children with a more aggressive “endotype 1” T1D version^76^, and the Manders overlap coefficient (MOC) used for quantification of ProINS/INS co-localization^76^ is contingent on confocal microscopy and not immediately comparable to the Jaccard index. Nevertheless, we note a similarity for individuals ≥13 years of age: a ProINS/INS MOC of ∼0.25 in Ctrl subjects that declines in T1DS cases^76^ seems to accord with a ProINS^hi^-INS Jaccard index of ∼25% in Ctrl donors that is significantly reduced in our T1DS and T1DL cohorts. **G.**, frequency distributions of islet sizes in PH (colored circles represent bin averages of all donors per respective group; linear log-log curve fits for all groups: R^2^ > 0.99). **H.,** left: distribution of islet sizes in the PH comprising all 12,200 individual islets stratified according to donor group and individual donors therein (within each donor group, donors are ordered according to increasing age; white bars: medians. Right: the violin plot summarizes donor group-specific islet size distributions (median/quartiles indicated) with statistical differences calculated using a mixed model (no significant differences recorded). **I.**, correlation of median islet size with donor age in PT and PH (goodness of linear regression fit and significance of slope deviation from zero are indicated). While AAb donors are ∼4 years older than other donors (*c.f.,* ***Table 1***), the ∼37% islet size increase in the PT of AAb *vs.* Ctrl donors (*c.f.,* ***Fig.2L***) exceeds an estimated age-associated islet size gain of ∼17% as shown here. **J.,** islet cell densities (geometric means of islet cell numbers per mm^2^ islet area). **K.,** islet sizes, diameters and volumes in PT *vs.* PH stratified according to T1D stage. To facilitate comparisons with the published literature, the panels feature both median and mean islet areas as determined from direct islet area measurements as well as median and mean islet diameters and volumes calculated from islet areas under the assumption of circularity = 1 and sphericity = 1 (*c.f.,* equations in ***Fig.1F***; although these assumptions constitute an oversimplification they are in line with similar calculations made in other reports). The values in the upper left panel corners indicate median or mean values for Ctrl donor islets in PT (dark blue) and PH (lighter blue) (the difference between median Ctrl PT *vs.* PH islet sizes, diameters and volumes remains non-significant even after outlier removal from the PH group). **L.,** frequencies and relative mass contributions of islets greater than average size in PT *vs.* PH stratified according to T1D stage. **M.,** average islet perimeters in PT and PH. **N.,** islet aspect ratio (Feret_min_ / Feret_max_ diameters), solidity (area / convex area) and circularity (4ν x area / perimeter^2^) in the PH; the binned histograms display donor group-specific islet circularity distributions. Due to notably weak CHGA and/or INS staining of PT but not PH sections from two donors (CHGA: Ctrl 6162; INS: Ctrl 6162, Aab 6450), the respective PT data are excluded in panels A, B and F. Statistical analyses were conducted with Student’s t-test (panels A, D left, K, L) or ordinary one-way ANOVA and Tukey’s multiple comparisons test (panels B, C, D right, E, F, J, M, N); unless noted otherwise, data are scatter and mean, and both t-tests and ANOVAs adopt the following convention: *p<0.05, **p<0.01, ***p<0.001, ****p<0.0001.

**Supplementary Figure S4. Correlations of islet size with islet properties & islet properties stratified according to UMAP cluster affiliation. A. – H.,** PT and PH islets ranging from 10^3^ - 10^5^μm^2^ area size from all donors within indicated T1D stage groups were allocated to 14 bins (variable islet size bins; 1,000-10,000μm^2^: 9 bins, 10,000-15,000μm^2^: 2 bins, 15,000-20,000μm^2^: 1 bin, 20,000-40,000μm^2^: 1 bin, 40,000-100,000μm^2^: 1 bin), and mean islet size within each bin is plotted against corresponding islet features (islet cell numbers, density, circularity, solidity and relative hormone staining areas). All curve fits describe exponential relations (one-phase association or decay) and goodness of fit (R^2^) is noted; in some cases, curve fitting was performed under exclusion of data points for very small islets as indicated by vertical dotted lines in the respective plots. Note that the data for circularity, solidity and islet cell density as well as endocrine, CHGA, ProINS^hi^, INS, IAPP, GCG and SST contents are the same as in *Fig.2P/Q* and shown here for clarification of curve fits. A. & B., Ctrl donors; C. & D., AAb donors; E. & F., T1DS donors; and G & H., T1DL donors. **I.,** left: histograms of islet size and ProINS area size distribution stratified according to T1D donor group; the vertical bar in ProINS histograms is set at 100μm^2^ corresponding to ∼1 endocrine cell, and values are percentages of islets with ProINS staining areas of ≥100μm^2^. Right: dot plot displaying islet area *vs.* ProINS area for individual islets color-coded according to T1D stage group. **J.,** histograms display cluster-stratified frequency distributions of combined PT and PH islets from all donor groups (Ctrl, AAb, T1DS, T1DL) according to indicated modalities (islet circularity and cellular density as well as relative endocrine and hormone staining areas for IAPP, ProGCG, SST, PPY); the adjacent heatmaps stratify the same parameters across T1D stage, individual donors (listed in order of increasing age within each group), islet cluster affiliation, and PT/PH regions. Note that not all donors have islets populating each cluster and we further omitted values if clusters contained <3 islets or <2 donors; missing and excluded values rendered in gray.

**Supplementary Figure S5. Islet properties across UMAP clusters and T1D stages. A. – D.,** scatter plots compare respective analytical parameters (scatter and mean) across UMAP clusters I, II, III, IV, V-A and V-BC, and are grouped according to T1D stage (top to bottom rows). A., islet shape descriptors for PT (Delaunay z-scores refer to a distance property of individual islets in relation to neighboring islets as detailed in ***Fig.6D***). B., relative islet endocrine hormone expression in the PT (“endocrine” in the first column of panel B refers to the union of all hormone staining areas). C., islet shape descriptors for the PH. D., relative islet endocrine hormone expression in the PH. **E. – H.,** the data featured in panels E. – H. are the same as in panels A. – D. but now all scatter plots compare respective analytical parameters across T1D disease stages (Ctrl, AAb, T1DS, T1DL) and are grouped according to UMAP cluster affiliation (top to bottom rows). E., islet shape descriptors for PT; F., relative islet endocrine hormone expression in the PT; G., islet shape descriptors for PH; and H., relative islet endocrine hormone expression in PH. Values on top of x-axes are the numbers of donors analyzed for each cluster and disease stage (“0” refers to absent values or data exclusion if <3 islets or <2 donors were represented in a given cluster, and cases with only two donors are highlighted in red to indicate the need for interpretative caution), and are applicable to all plots in the same panel row. Features specifically called out in the main narrative are highlighted with gray background for easier identification. Statistical analyses were conducted with ordinary one-way ANOVA and Tukey’s multiple comparisons test adhering to the following convention: *p<0.05, **p<0.01, ***p<0.001, ****p<0.0001.

**Supplementary Figure S6. UMAP islet cluster features, islet properties & CD45 cell infiltration.** Note that in panels A, B, C and G not all donors have islets populating all clusters, and in addition we excluded values if <3 islets or <2 donors were represented in a cluster; the resultant numbers of donors analyzed for each cluster are featured on top of the respective x-axes with absent values indicated by “0” and the presence of only two donors highlighted in red font to indicate the need for interpretative caution. **A.,** combined islet size distribution in UMAP cluster map (left), and cluster-level comparison of islet area z-scores in PT *vs.* PH across T1D disease stages (right). **B.,** combined islet circularity distribution in UMAP cluster map, and cluster-level comparison of islet circularity z-scores in PT *vs.* PH across T1D disease stages (right). **C.,** cluster-level islet cell densities in PH (all clusters I – III *vs.* IV – V-BC: p<0.0001). **D.,** relative magnitude of UMAP islet subclusters I-A and I-B in PT and PH across T1D disease stages. To ascertain the collapse of cluster I from Ctrl to AAb stage in more detail, we considered its subclusters I-A/B/C where the main difference pertains to the presence (I-A) or absence (I-B) of delta cells (*c.f.,* ***Fig.3C***) (the minor subcluster I-C is only found in the PH and contains islets with beta, gamma and delta but not alpha cells; *c.f.,* ***Fig.3A/C***). Compared to Ctrl donors, AAb donors present with a 45% reduction in PT clusters I-A/B and PH cluster I-A as well as a lesser 15% decrease in PH cluster I-B. **E.,** relative magnitude of UMAP islet subclusters IV-A – D in the PH across T1D disease stages **F.,** relative magnitude of UMAP islet clusters I – V in PT *vs.* PH regions across T1D disease stages (*c.f.,* cluster size comparisons in ***Fig.4C/D/F***). In aggregate, the consideration of donor-specific islet cluster size distributions permits the identification and characterization of several outliers: Ctrl 6454 presents with an “AAb-like” cluster II dominance that would appear consistent with their unusually high beta and alpha cell mass (*c.f., **Fig.S2K***); AAb 6310, despite being only single AAb^+^ at time of death and with a low genetic HLA risk, exhibits a pattern comparable to donors with multiple AAbs and thus may have been at greater T1D risk than clinically apparent; AAb 6450 has a profile resembling T1DS donors, moderate HLA risk and might have been poised to develop clinical disease in the near future; and T1DS 6405 resembles AAb donors and is distinct from other T1DS donors on account of severe obesity, excessive pancreas weight, elevated C-peptide and comparatively lower Hb1Ac despite high HLA risk (*c.f., **Fig.S2A-D***). **G.,** scatter plot summaries depicting peri- and intra-islet CD45 cell frequencies in PH clusters II – V-A. **H.,** peri-islet : intra-islet CD45 frequency ratios in clusters II and V-A of PT and PH. **I.,** correlations of total islet-associated CD45 cell numbers, CD45 cell frequencies and islet size in cluster II (all donors, PT/PH combined). CD45 number binning was performed in one CD45 cell increments, CD45 percentage binning in 0.5% increments, and islet size binning employed 1,000μm^2^ increments; curve fits correspond to exponential associations and goodness of fit (R^2^) is indicated (the vertical or horizontal red broken lines indicate the insulitis threshold of ≥15 CD45 cells). “CD45 # binning” documents a straightforward exponential association with corresponding CD45 cell frequencies that is particularly good for ∼1-8 CD45 cells; similarly, “CD45 % binning” reveals a quasi-linear association that again is especially tight for islets with ∼1-9 CD45 cells. However, these relations are not strictly reciprocal since they yield somewhat different CD45 cell frequencies corresponding to the insulitis threshold (“CD45 # binning”: 5.7%; “CD45 % binning”: 8.5%; also note that the curve fit for CD45 frequency bins *vs.* islet size [“CD45 % binning”, right plot] demonstrates a brief exponential decline followed by a plateau starting at CD45 cell frequencies of ∼1.5% - 2.0%). Importantly, “islet size binning” yields a positive correlation of islet size with CD45 numbers but a negative correlation with CD45 frequencies since small denominators inflate relative frequencies of CD45 cells even if their numbers are low. **J.,** combined PT and PH cluster II (top) and cluster I (bottom) islets stratified according to Ctrl, AAb or T1DS stage were “size-binned” for comparison of associated CD45 cell frequencies, shape descriptors and relative hormone expression areas (Ctrl cluster I: 5,706 islets, cluster II: 1,750 islets; AAb cluster I: 2,279 islets, cluster II: 2,426 islets; T1DS cluster I: 640 islets, cluster II: 745 islets; variable islet size bins [1,000-10,000μm^2^: 9 bins, 10,000-15,000μm^2^: 2 bins, 15,000-20,000μm^2^: 1 bin, 20,000-40,000μm^2^: 1 bin, 40,000-100,000μm^2^: 1 bin]; each data point is the mean of all cluster II or cluster I islets within respective size bins; where applicable, curve fits represent exponential associations). Statistical analyses were conducted with ordinary one-way ANOVA and Tukey’s multiple comparisons test (panels C-E, G and H) or paired Student’s t-test (panels A, B and F) with *p<0.05, **p<0.01, ***p <0.001, ****p<0.0001.

**Supplementary Figure 7. Spatial distribution of UMAP-clustered islet subsets across pancreatic tissue sections and T1D disease stages (I). A.,** modified Ripley’s K function analyses were conducted as detailed in *Fig.6A-C* and Methods. One-way ANOVA tests were performed for each distance bin and are significant for the range of 1.5-8×10^3^μm in PT, and 0.6-10×10^3^μm in PH (PH 6380 excluded due to very scarce islets, *c.f. Fig.S8A*); area under Ripley curve calculations displayed here only pertain to the above distance ranges. **B.,** islet distribution in each tissue section can also be described using fractal geometry (see Methods): boundary-adjusted islet counts found at each search radius were fit to a power function; each point here is the exponent calculated for a single slide; and a random distribution of islets would lead to a value of 2 while smaller values indicate section-wide clustering (statistical analyses conducted with ordinary one-way ANOVA and Tukey’s multiple comparisons test with *p<0.05 and **p<0.01). **C.,** mean Delaunay distances between islets across and distribution of mean Delaunay distances according to T1D stage. **D.,** Delaunay distance distributions of islets grouped according to cluster affiliation and disease stage in PT and PH; values in each histogram indicate the corresponding mean Delaunay distances. Note that a marked rise of Delaunay distances for cluster I islets in AAb donors further increases with T1D onset; a similar pattern is observed for cluster II islets and cluster III islets in the PH; and Delaunay distances for cluster V islets in the PT but not PH expand from AAb to T1DL stage. A notable exception pertains to cluster IV islets in the PH which exhibit an overall greater proximity that appears to grow even closer with T1D progression; this likely results from a combination of unique regionalization, an imperviousness to T1D pathological processes due to constitutive lack of beta cells, and a merging with former cluster III islets rendered beta cell-deficient (*c.f., Fig.4E/G*). **E.,** heatmaps stratifying Delaunay distances (left) and Delaunay area z-scores (right) across T1D stage, individual donors, islet cluster affiliation, and PT and PH regions (missing/excluded values in gray); Delaunay area z-scores are also featured in the summary *Fig.S5*. **F.,** contour plots are gated on combined islets from all donor groups and display islet size against mean Delaunay distance; quadrant markers distinguish larger islets (>10,000μm^2^) and greater Delaunay distances (>1,000μm), and values are percentages of islets in each quadrant. Note that both larger and smaller islets are subject to the disease-associated increase of Delaunay distances. **G.,** Delaunay distances for insulitic and other islets (histograms gated on combined PT and PH islets from AAb *vs.* T1DS donors; values are the respective fractions of islets at >1,250μm Delaunay distance). Both non-insulitic and insulitic islets display essentially similar neighborhood relations with small subsets (∼6% in AAb, ∼11% in T1DS) that are more “isolated” at a distance of >1,250μm. H., *in situ* projection of cluster color-coded islets onto a representative PH section. **I.,** approximation of islet cluster stratification with only INS, GCG and CD45 stains; in the absence of additional PPY stains, cluster III islets in the PH cannot be distinguished from GCG-negative cluster I or II islets, and cluster IV islets remain “invisible” (the callout for panel H is found in the Conclusions section).

**Supplementary Figure 8. Spatial distribution of UMAP-clustered islet subsets across pancreatic tissue sections and T1D disease stages (II).** Pancreatic tissue section outlines (regions, T1D stage and nPOD donor IDs are indicated) populated with color-coded islets; for facilitated visual presentation, all islets are rendered circular and enlarged with relative size differences preserved. **A.,** islets color-coded according to UMAP cluster affiliation. **B.,** islets color-coded according to the numbers (0-15+) of associated CD45 cells. Distinctive histopathological properties of three “outlier” cases are readily visualized in this data display: AAb 6450 with a histopathological presentation that appears “T1DS-like”; AAb 6310 who despite single AAb positivity features islet subset distributions similar to AAb donors with ≥2 AAbs (*c.f., Fig.4B*); and T1DS 6405, the donor with severe obesity and better glucose handling (*c.f., Fig.S2B/C*) whose pancreas histology resembles AAb or even Ctrl donors.

## METHODS

### RESOURCE AVAILABILITY

#### Lead Contact

Requests for further information and resources should be directed to and will be fulfilled by the lead contact, Dirk Homann.

#### Materials availability

Not applicable.

#### Data and code availability

Code used for image analysis using QuPath and MATLAB is accessible on GitHub: https://github.com/saramcardle/Image-Analysis-Scripts/tree/master/MICSSS_Pancreas. All islet measurements are featured in ***Table S2***. Any additional information required to reanalyze the data reported in this manuscript is available from the lead contact upon request.

### EXPERIMENTAL SPECIMENS DETAIL

Formalin-fixed, paraffin-embedded (FFPE) tissue sections (5μm) from pancreatic tail and head regions of organ donor pancreata were provided by the Network for Pancreatic Organ Donors with Diabetes (nPOD) and comprise 25 donors allocated to four donor groups: 7 non-diabetic control donors (Ctrl), 6 autoantibody-positive (AAb) donors, 8 donors with short duration of clinical type 1 diabetes (T1DS, <2 years), and 4 donors with longer duration of clinical T1D (T1DL, 8-11 years); donor matching across the 4 groups was performed on age and gender with additional matching for demographic (ethnicity) and clinical (body mass index/BMI) parameters where possible; further details including demographic and clinical metadata are provided in ***Table S1***. To establish and optimize staining protocols, additional pancreatic and splenic FFPE tissue sections were provided by the nPOD consortium and the ISMMS Biorepository and Pathology CoRE.

### METHOD DETAILS

#### Immunohistochemistry (MICSSS)

FFPE tissue sections (5 µm) were sequentially stained for eight islet hormones and CD45^+^ immune cells adjusting the ochemical consecutive staining on single slide (MICSSS) method^41,42^. Iterative staining order was empirically determined for pancreatic tissues to account for differential antigen sensitivity to deterioration during consecutive MICSSS cycles. Briefly, slides were “baked” overnight (o/n) at 37°C to ensure tissue adherence to slides in subsequent staining rounds. Following deparaffinization in histology-grade Xylene (Fisher Scientific), sections were rehydrated by immersing them in a series of graded ethanol solutions (histology-grade, Fisher Scientific) at decreasing concentrations (3×100%, 90%, 70%, and 50%) down to distilled water (5 min each) prior to heat-induced epitope retrieval (HIER) at pH6 (Citrate Buffer; ThermoFisher Scientific) in a 95°C water bath for 30 min. Tissue sections were cooled down to room temperature, endogenous hydrogen peroxidase activity was quenched by incubation with PeroxAbolish solution (Biocare Medical) for 10 min and slides were subsequently washed in Tris Buffered Saline (TBS, Cell Signaling; 2x 5 min). Non-specific background due to Fc receptor binding was blocked by incubation with DAKO serum-free protein block (Agilent) for 15 min, endogenous biotin was blocked using the DAKO Biotin-Blocking System (Agilent) according to manufacturer instructions before addition of primary antibodies listed as specified in ***Table S3***. Target antigens were revealed after incubation with biotinylated F(ab’)_2_ donkey-raised secondary antibodies with minimal cross-reactivity (Jackson ImmunoResearch; 30 min RT), horseradish peroxidase (HRP)-conjugated streptavidin (DAKO, Agilent; 30 min), and ImmPACT AMEC Red substrate (Vector Laboratories) as per vendor’s instructions. Tissue sections were counterstained with Harris modified hematoxylin solution (Sigma), mounted with an aqueous mounting medium (DAKO Glyercgel, Agilent), whole-slide images were acquired at 40x on a NanoZoomer S60 Digital Slide Scanner (Hamamatsu) and exported as .ndpi files (for some preparatory experiments, images were acquired at 20x using a Pannoramic 250 Flash II Digital Scanner [3DHistech]; *c.f.* ***Fig.1B***). Slides were stored protected from light at 4°C until further staining. For sequential staining of subsequent targets in MICSSS cycles #2-9, cover slips were carefully removed in hot water (∼50°C) and tissue sections were rinsed in distilled water, destained/dehydrated by immersing them in ethanol solutions at increasing concentration (50, 70, 100%; 3 min each) and xylene (3×2 min), and subsequently rehydrated through a graded ethanol series and distilled water as detailed before. Antigen retrieval at pH6 was performed for 10 min as specified above removing hematoxylin staining, endogenous and streptavidin-associated (from previous rounds of staining) hydrogen peroxidase activity, non-specific background, as well as endogenous and secondary antibody-mediated (from previous rounds of staining) biotin were blocked using PeroxAbolish, DAKO serum-free protein block, and DAKO Biotin-blocking system as above. To enable probing for target antigens using primary antibodies raised in the same species, tissue sections were incubated with 5% mouse or rabbit IgG (serum; Jackson ImmunoResearch) (depending on the primary antibody used in the immediate prior staining cycle) for 1h at RT followed by incubation with donkey-raised AffiniPure Fab Fragments (Jackson ImmunoResearch) against the previously used primary antibody species (200 µg/mL) at 4°C o/n. Subsequent primary antibody staining, target revelation, and image acquisition were conducted as described above.

#### QuPath analysis pipeline

##### Image alignment

Images were imported into QuPath version 0.2.3^37^, with each set of 9 images of a single slide bundled into an individual project. Using the CD45 image as the “base” image, the other 8 project images were each aligned to it using a custom groovy script based on QuPath’s Interactive Image Alignment function, incorporating affine transforms at 5µm resolution. A threshold pixel classifier was used to detect the entire tissue area on the CD45 image, which had been acquired last and therefore exhibited all of the accumulated tissue damage from the iterative MICSSS staining process. The pancreas tissue was then transferred to each image in the project, utilizing the pre-calculated affine transform matrix.

##### Islet detection

Further image processing was performed with QuPath version 0.4.3 or 0.5.0. Stain separation vectors were optimized for each hormone staining round and consistently applied to every image with that antibody to spectrally unmix the AMEC chromogen from hematoxylin. CHGA was expected to mark all islets yet staining was notably faint in some PPY^+^ and small GCG^-^ islets; we therefore delineated islets with a combination of six hormone stains (CHGA, ProINS, INS, GCG, SST and PPY; ProGCG and IAPP proved redundant for this task and were therefore not included to reduce computational complexity and time). On these six stains, we segmented objects with a low-resolution pixel classifier (0.88μm), thresholding the AMEC channel to find all stained regions with an area of at least 50μm^2^. All objects were transferred to their related CD45 base image and then merged. The large resulting annotation was split into individual objects. Holes were filled in to create contiguous objects, and then all objects less than 1,000μm^2^ were removed. Additionally, objects that were within 10μm of the tissue border were removed to prevent errors from incomplete islet capture and edge staining artifacts. The proto-islets were converted to detections and redistributed to all nine stains, where their overall AMEC staining intensity was measured (mean, standard deviation, min, max, and Haralick features for texture). The measurements from each object were gathered, along with shape features, and used to train a machine-learning object classifier to remove artifacts due to dust, non-specific staining, tissue damage, or loss of focus. The classifier was trained on multiple representative images from all four donor groups. We then measured and calculated shape descriptors for each final islet, and we performed Delaunay clustering on each slide with a maximum search radius of 4,000μm (1 of 24,578 islets had no neighbors within 4mm; this islet was assigned a Delaunay distance of 4mm and a Delaunay area of ν x 16mm^2^).

##### Hormone staining areas & islet/endocrine cell type mass

Islet boundaries were transferred to each of the eight hormone-stained images and a high-resolution (0.22μm) pixel thresholder was applied to find regions of positive staining. ProINS was the first stain in the series and therefore allowed for reliable differentiation of darker staining, lighter staining, and background. As CHGA is expected to be in nearly all endocrine cells, we captured all staining areas including lighter stained sections. For all other hormones, we only captured darker staining areas to avoid inclusion of background signal. The detections representing the stained area per islet were all returned to the base image and the positive area in each islet was recorded as a percentage of total islet size. For every pair of hormone stains (56 pairs), the Java JTS topography suite was used to calculate the intersection and the union of the stained regions. These areas were subsequently used to calculate Jaccard indices and relative areas of double-positive regions per islet. Additionally, we merged all hormone stains to find the total endocrine area per islet. Lastly, islet and endocrine cell type mass was calculated by multiplying relative fractions of islet areas or specific hormone staining areas with regional pancreas weights (PT or PH) (***Table S1***).

##### Immune cells

Islet boundaries were expanded by 20μm to mark the peri-islet region, using a watershed algorithm to ensure that the peri-islet boundaries of neighboring islets did not overlap. Within these regions, nuclei were detected *via* the QuPath implementation of Stardist^48^, with 1μm expansion for the cytoplasm. An object classifier was trained to detect CD45^+^ immune cells. As the last stain in the MICSSS series, the CD45 image had the highest diffuse background staining within islets which furthermore varied between samples. To resolve this issue, the classifier used measurements of CD45 staining intensity in the nuclear, cytoplasmic, and membrane compartments, as well the smoothed average of cells within a 100μm neighborhood, and the difference between the cell’s intensity and that of a 10μm circular tile, including extracellular regions. After classification, the number and frequency of CD45^+^ cells inside the islet and peri-islet region were calculated.

##### CytoMAP analysis

Islet measurements were exported to a .csv file and used for dimensionality reduction and clustering in the MATLAB implementation of CytoMAP^91^. The data input into UMAP^71^ for dimensionality reduction included: the area of eight hormone stains, the total (union) endocrine stain area, islet-associated CD45^+^ cell count, circularity, mean Delaunay area, and islet area. The total area, Delaunay area, and raw hormone areas (μm^2^) appeared log-normally distributed while the hormone staining areas as a percentage of islet areas were far from a normal distribution. Therefore, we used the log-transformed raw areas consistently. To balance the varying inputs, we calculated the z-scores for all of them. Pre-processing was performed in MATLAB. UMAP was run with the following parameters: n_neighbors = 50, min_dist = 0.1, n_epochs = 1000. Two rounds of DBScan clustering were performed on the UMAP output. First, to distinguish larger clusters, we used a high epsilon value (0.5) with a minimum number of 50 points. This yielded six clusters that largely corresponded to the visible clustering of the UMAP scatter plot (clusters 5 and 6 are in close proximity and were named V-A and V-BC). Second, to distinguish subclusters that were largely delineated by SST and GCG, we used a low epsilon (0.2) and a low minimum number of points (20). This yielded 16 clusters with some unclustered points. We used these results to divide the six major clusters (I, II, III, IV, V-A, V-BC) into subclusters for deeper analysis. The islet cluster assignment was exported from CytoMAP into a .csv file and reuploaded into QuPath as a detection measurement using custom scripts. The cluster ID was converted into a class and then class-specific Delaunay clustering with a maximum search radius of 8mm was performed.

##### Spatial analysis

For each slide, the automatically created tissue outline was manually edited to remove peripheral areas of adipose and connective tissue and fill in regions of missing tissue to determine the pancreas parenchymal area. This was exported from QuPath as a .geojson file and then imported into MATLAB along with the islet centroid locations. To calculate spatial distribution patterns in each slide, we used a modified Ripley’s K function^92,93^ that incorporates the tissue boundary and the total pancreas density. Circles are placed at each islet with varying radii (400 - 10,000μm); looping through each islet, the number of other islets within that circle is counted; and the fraction of the circle area that lies inside the tissue boundary is used as a weighting factor for the number of points found. The adjusted number of points found at each distance is averaged across all islets in a slide. For Ripley analysis, this count is normalized to the number of points expected if islets were randomly distributed in the pancreas area - the average density multiplied by the area of the circle. If, at a given radius, the islets had on average the expected number of neighboring islets, the modified K value would be 1; however, if there were twice as many islets within a radius as expected, the modified K value would be 2. This allows for a comparison of spatial islet aggregation even if the pancreatic tissue sections have different sizes, islet densities, and complexities. These data were further used for fractal spatial analysis following the method of Jo *et al.*^85^. For each slide, search radii and boundary-adjusted islet counts were plotted on a log-log graph and the slope of the best-fit line was calculated, excluding radii ≤600µm; the slopes, or fractal dimensions, of each slide were compared to demonstrate average changes in islet distribution with disease progression.

##### Data visualization

To ease data visualization and figure creation, the large matrix of islet measurements, including single and double hormone areas, locations, shape descriptors, spatial measurements and UMAP clusters were converted to .fcs files for visualization in Flowjo 10.10.0 (BDBiosciences) using the writeFCS function^94^. In some cases, contrast and brightness were adjusted for entire brightfield images using Adobe Photoshop, and scale bars were added in Fiji/imageJ. Pseudofluorescence images were generated in QuPath using the spectrally separated AMEC channels per stain and the previously calculated affine transforms.

#### HLA haplotype T1D risk classification

HLA class I and class II haplotype data were classified for T1D risk according to published data and are summarized in ***Table S1***. HLA class II DR-DQ genotypes were binned based on T1D risk, high, moderate, neutral or protective, determined from individuals of European ancestry^52^. Bins were assigned numerical values for the purposes of this study, high risk (+2), moderate risk (+1), neutral (0), or protective (−1). Alternatively, HLA class II risk was classified using published odds ratios (OR) calculated for individuals of European (DR-DQ haplotype combinations^53^), African (individual DR-DQ haplotypes^54,55^), or admixed Hispanic/Latino descent (individual DR-DQ haplotypes^55^). HLA class I risk was classified for HLA-A and -B genotypes using OR calculated from individuals of European descent^95^.

### QUANTIFICATION AND STATISTICAL ANALYSIS

Data analysis and graphical representation were performed in GraphPad Prism 9 and 10 (GraphPad Software). Normal distribution was determined by D’Agostino-Pearson test. Statistical significance was assessed by unpaired or paired Student’s t tests or non-parametric Mann-Whitney U test as applicable for comparison between two groups; by one-way ANOVA with Tukey’s multiple comparisons post hoc testing for analysis of more than two groups with normally distributed values; or by one sample t test using a hypothetical mean as indicated. Analyses of islet sizes were conducted using mixed effect models with subjects as random effects specifying a compound-symmetry correlation matrix in the model, and orthogonal contrasts were used for pair-wise comparison of the four donor groups. Summary data are displayed as scatter plots with mean, violin plots with median and quartiles, bar diagrams (mean±SE), frequency distributions (mean or median with interquartile range as indicated), or correlation plots adopting the following convention: *p<0.05, **p<0.01, ***p <0.001, ****p<0.0001; ns or no symbol, non-significant.

Missing and excluded data: for the purpose of the present study, pancreatic islets are defined as endocrine objects ≥1,000μm^2^ (∼10 cells, ∼36 μm diameter); accordingly, we excluded smaller endocrine structures and single cells from all of our analyses. Information about HLA haplotypes and HbA1c values was available for most but not all donors (***Table S1 & Fig.S2A/C***). Subcluster IV-E (***Fig.3A***) is sample-biased and was not further considered. In some analyses of UMAP cluster-stratified islets, not all donors have islets present in all clusters; in addition, we excluded values if <3 islets or <2 donors were represented in a cluster. The resultant numbers of donors analyzed for each cluster are featured on top of respective x-axes in ***Figs.6I/J, S5 & S6A-C/G/H*** with absent values indicated by “0” in red font and the presence of only two donors also highlighted in red font to indicate the need for interpretative caution. CHGA and INS staining of Ctrl 6162 PT tissue sections and INS staining of AAb 6450 PT sections was notably weak; in the absence of CHGA and INS staining irregularities in the corresponding PH sections as well as normal ProINS and IAPP staining in both PT and PH sections, we attribute this observation to a technical staining issue and therefore excluded the respective PT CHGA and INS data from ***Figs.2D, 3D, S1C, S2I-K, S3A/B/F & S5B/F***. (***Figs.2B/C & S2K***), and PH data from T1DS 6380 was excluded from modified Ripley’s K analyses due to very scarce islets (***Fig.6C* & *S8A***).

