## Supplemental Tables 1 and 3, Supplemental Figures S1 - S8 for "Integrated histopathology of the human pancreas throughout stages of type 1 diabetes progression"

Table S1

**Table S1. Pancreas specimen information & donor metadata**

| case / tissue information | | | | | demographic & clinical parameters | | | | | | | | | | | pancreas weight (g) | | | | | insulitis* |
|---|---|---|---|---|---|---|---|---|---|---|---|---|---|---|---|---|---|---|---|---|---|
| case ID & block # | pancreas head / tail | ID (other) | slide type | donor type / T1D stage | gender | ethnicity | age (years) | age at T1D onset (years) | T1D duration (years) | body weight (kg) | BMI | C-peptide (ng/ml) | HbA1c (%) | # of AAb (type of AAb) | cause of death | total | H (head) | B (body) | T (tail) | sum H, B & T | nPOD assessment |
| **7 non-diabetic control cases (Ctrl)** | | | | | | | | | | | | | | | | | | | | | |
| 6382-02<br>6382-02 | H<br>T | 20620<br>20621 | paraffin<br>paraffin | Control<br>(non-diabetic) | F | CEU | 4.7 | n/a | n/a | 19.7 | 17.2 | 5.21 | 5.30 | 0 (negative) | head trauma | 17.02 | 6.28 | 5.30 | 5.44 | 17.02 | no |
| 6278-02<br>6278-02 | H<br>T | 20610<br>20611 | paraffin<br>paraffin | Control<br>(non-diabetic) | F | AA | 12.0 | n/a | n/a | 52.4 | 21.3 | 4.54 | 6.30 | 0 (negative) | anoxia | 33.60 | 11.90 | 13.50 | 8.20 | 33.6 | no |
| 6386-02B<br>6386-02A | H<br>T | 20623<br>20622 | paraffin<br>paraffin | Control<br>(non-diabetic) | M | CEU | 14.0 | n/a | n/a | 71.5 | 23.9 | 1.12 | 5.60 | 0 (negative) | head trauma | 61.36 | 18.30 | 18.19 | 24.87 | 61.36 | no |
| 6389-02A<br>6389-04 | H<br>T | 20624<br>20625 | paraffin<br>paraffin | Control<br>(non-diabetic) | M | CEU | 18.6 | n/a | n/a | 70.0 | 20.9 | 7.22 | 5.10 | 0 (negative) | head trauma | 65.85 | 20.65 | 22.44 | 22.76 | 65.85 | no |
| 6162-02<br>6162-01 | H<br>T | 20596<br>20597 | paraffin<br>paraffin | Control<br>(non-diabetic) | M | AA | 22.7 | n/a | n/a | 102.0 | 28.9 | 7.61 | n.d. | 0 (negative) | head trauma | 81.50 | 19.20 | 26.40 | 31.50 | 77.1<br>(Δ-4.4 = 5.4%) | no |
| 6401-02A<br>6401-02 | H<br>T | 20628<br>20629 | paraffin<br>paraffin | Control<br>(non-diabetic) | F | AMR | 25.1 | n/a | n/a | 85.3 | 31.3 | 12.81 | 5.80 | 0 (negative) | head trauma | 71.29 | 23.72 | 21.20 | 26.37 | 71.29 | no |
| 6454-02B<br>6454-02 | H<br>T | 20640<br>20641 | paraffin<br>paraffin | Control<br>(non-diabetic) | M | CEU | 25.2 | n/a | n/a | 89.0 | 26.6 | 4.57 | 4.70 | 0 (negative) | anoxia | 55.67 | 24.84 | 17.06 | 13.77 | 55.67 | no |
| **6 auto-ab+ cases (AAb)** | | | | | | | | | | | | | | | | | | | | | |
| 6424-02A<br>6424-04B | H<br>T | 20634<br>20635 | paraffin<br>paraffin | AutoAb+<br>(stage 1/2) | M | CEU | 17.7 | n/a | n/a | 176.0 | 51.4 | 6.97 | 5.80 | 2 (GADA, mIAA) | head trauma | 111.07 | 32.89 | 43.52 | 34.66 | 111.07 | no |
| 6197-06<br>6197-04 | H<br>T | 20600<br>20643 | paraffin<br>paraffin | AutoAb+<br>(stage 1/2) | M | AA | 22.0 | n/a | n/a | 89.0 | 28.2 | 17.48 | 5.50 | 2 (GADA, IA2A) | head trauma | 73.30 | 14.20 | 38.80 | 20.30 | 73.3 | yes,<br>"rare" |
| 6450-02C<br>6450-04A | H<br>T | 20639<br>20638 | paraffin<br>paraffin | AutoAb+<br>(stage 1/2) | F | CEU | 22.0 | n/a | n/a | 66.0 | 24.4 | 5.47 | 5.70 | 2 (GADA, ZnT8A) | anoxia | 54.18 | 21.84 | 16.18 | 16.16 | 54.18 | yes |
| 6429-02A<br>6429-02 | H<br>T | 20636<br>20637 | paraffin<br>paraffin | AutoAb+<br>(stage 1/2) | M | AA | 22.1 | n/a | n/a | 62.1 | 19.6 | 2.25 | 5.50 | 2 (GADA, mIAA) | head trauma | 70.66 | 26.85 | 22.01 | 21.80 | 70.66 | no |
| 6267-02A<br>6267-01 | H<br>T | 20609<br>20642 | paraffin<br>paraffin | AutoAb+<br>(stage 1/2) | F | CEU | 23.0 | n/a | n/a | 64.0 | 23.5 | 16.59 | 5.00 | 2 (GADA, IA2A) | anoxia | 47.99 | 15.30 | 15.45 | 17.24 | 47.99 | yes |
| 6310-02B<br>6310-02B | H<br>T | 20612<br>20613 | paraffin<br>paraffin | AutoAb+<br>(stage "0") | F | AMR | 28.0 | n/a | n/a | 63.0 | 22.4 | 10.54 | n.d. | 1 (GADA) | anoxia | 82.93 | 26.39 | 27.64 | 28.90 | 82.93 | yes,<br>"low-grade" |
| **8 T1D short cases (T1DS)** | | | | | | | | | | | | | | | | | | | | | |
| 6209-01<br>6209-02 | H<br>T | 20601<br>20602 | paraffin<br>paraffin | T1D short<br>(stage 3) | F | CEU | 5.0 | 4.75 | 0.25 | 15.0 | 15.9 | 0.10 | n.d. | 3 (IA2A, ZnT8, mIAA) | cerebral edema, DKA | 12.50 | 4.70 | 4.50 | 3.30 | 12.5 | yes |
| 6380-04A<br>6380-02 | H<br>T | 20619<br>20618 | paraffin<br>paraffin | T1D short<br>(stage 3) | F | AA | 11.6 | 11.60 | 0.00 | 30.8 | 14.6 | 0.22 | 13.50 | 0 (negative) | cerebral edema, DKA | 34.26 | 10.54 | 14.29 | 9.43 | 34.26 | "possibly" |
| 6371-02<br>6371-02 | H<br>T | 20617<br>20616 | paraffin<br>paraffin | T1D short<br>(stage 3) | F | CEU | 12.5 | 10.50 | 2.00 | 40.0 | 16.6 | 0.11 | 9.50 | 4 (GADA, IA2A, mIAA, ZnT8A) | cerebral edema | 22.76 | 6.28 | 7.07 | 9.41 | 22.76 | yes |
| 6228-02*<br>6228-01* | H<br>T | 20603<br>20604 | paraffin<br>paraffin | T1D short<br>(stage 3) | M | CEU | 13.0 | 13.00 | 0.00 | 45.0 | 17.4 | 0.10 | 13.30 | 3 (GADA, IA2A, ZnT8) | anoxia | 30.35 | 11.15 | 10.75 | 8.45 | 30.35 | yes |
| 6396-02B<br>6396-04 | H<br>T | 20627<br>20626 | paraffin<br>paraffin | T1D short<br>(stage 3) | F | CEU | 17.1 | 15.10 | 2.00 | 71.6 | 22.6 | 0.06 | 13.40 | 0 (negative) | cerebral edema, DKA | 45.00 | 14.60 | 13.20 | 17.20 | 45 | yes |
| 6247-01<br>6247-03 | H<br>T | 20606<br>20605 | paraffin<br>paraffin | T1D short<br>(stage 3) | M | CEU | 24.0 | 23.40 | 0.60 | 81.2 | 24.3 | 0.47 | n.d. | 1 (mIAA) | head trauma | 59.77 | 20.80 | 21.41 | 17.56 | 59.77 | yes |
| 6362-04A<br>6362-02 | H<br>T | 20615<br>20614 | paraffin<br>paraffin | T1D short<br>(stage 3) | M | CEU | 24.9 | 24.90 | 0.00 | 90.0 | 28.5 | 0.38 | 10.00 | 1 (GADA) | head trauma | 53.26 | 13.13 | 17.17 | 22.96 | 53.26 | yes |
| 6405-02A<br>6405-04 | H<br>T | 20631<br>20630 | paraffin<br>paraffin | T1D short<br>(stage 3) | F | AMR | 29.1 | 28.50 | 0.60 | 108.8 | 42.5 | 1.84 | 7.00 | 3 (GADA, IA-2A, ZnT8A) | cerebro-vascular stroke | 94.27 | 34.29 | 29.75 | 30.23 | 94.27 | yes |
| **4 T1D long cases (T1DL)** | | | | | | | | | | | | | | | | | | | | | |
| 6264-02<br>6264-01 | H<br>T | 20607<br>20608 | paraffin<br>paraffin | T1D long<br>(stage 3) | F | CEU | 12.0 | 3.00 | 9.00 | 34.0 | 22.0 | <0.5 | 8.90 | 0 (negative) | DKA | 20.35 | 8.92 | 7.10 | 4.33 | 20.35 | yes<br>(PanBody 03) |
| 6089-06<br>6089-03 | H<br>T | 20595<br>20594 | paraffin<br>paraffin | T1D long<br>(stage 3) | M | CEU | 14.3 | 6.30 | 8.00 | 69.0 | 26.0 | <0.5 | 10.40 | 1 (mIAA) | anoxia | 56.80 | 12.10 | 15.90 | 25.70 | 53.7<br>(Δ-3.1 = 5.5%) | no |
| 6418-02B<br>6418-01 | H<br>T | 20632<br>20633 | paraffin<br>paraffin | T1D long<br>(stage 3) | M | CEU | 24.9 | 13.90 | 11.00 | 65.1 | 26.4 | <0.02 | 7.50 | 4 (GADA, IA2A, mIAA, ZnT8A) | anoxia | 28.59 | 12.38 | 7.70 | 8.51 | 28.59 | no |
| 6180-02<br>6180-01 | H<br>T | 20598<br>20599 | paraffin<br>paraffin | T1D long<br>(stage 3) | M | CEU | 27.1 | 16.10 | 11.00 | 82.0 | 25.9 | <0.05 | n.d. | 4 (GADA, IA2A, mIAA, ZnT8A) | head trauma | 36.70 | 12.50 | 11.90 | 12.30 | 36.7 | no |

| case ID & block # | HLA haplotypes & T1D risk | | | | | | | | | |
|---|---|---|---|---|---|---|---|---|---|---|
| | HLA-DRB1 | HLA-DQB1 | HLA-DQA1 | PMID 23539116 (CEU) | PMID 30655379 (CEU) | PMIDs 38453145 & 30659077 (AFR, AMR) | HLA-A | PMID 18004301 (CEU) | HLA-B | PMID 18004301 (CEU) |
| 6382-02 / 6382-02 | 03:01/11:01 | 02:01/03:01 | 05:01/05:05 | protective (-1) | 1.31 | na | 11:01/33:05 | 0.52/0.24 | 65/35 | 0.86 |
| 6278-02 / 6278-02 | 11:04/12:01 | 03:01/05:02 | 03:01/05:01 | protective (-1) | 1 | 0.17/na | 23:01/68:02 | 0.71/0.99 | 45/71 | 0.68 |
| 6386-02B / 6386-02A | 04:04/13:01 | 03:02/06:03 | 01:03/03:01 | neutral (0) | 5.53 | na | 02:01/03:01 | 1/1.13 | 18/44 | 1.83 |
| 6389-02A / 6389-04 | 04:01 | 03:01/06:02 | 01:02/03:03 | protective (-1) | 0.15 | na | 02:01/02:01 | 1/1 | 07/44 | 1.25/1 |
| 6162-02 / 6162-01 | 13:03/16:02 | 03:01/05:02 | 01:02/05:01 | neutral (0) | 1 | 0.36/0.93 | 25:01/68:02 | 1.04/0.99 | 57/58 | 0.55/0.53 |
| 6401-02A / 6401-02 | 07:01/13:01 | 02:02/06:03 | 02:01/01:03 | protective (-1) | <1 (no data) | 0.5/na | 31:01/33:03 | 0.94/0.24 | 14:02/51:01 | 0.66/0.93 |
| 6454-02B / 6454-02 | na | na | na | na | na | na | na | na | na | na |
| 6424-02A / 6424-04B | 03:01/04:01 | 02:01/03:02 | 05:01/03:01 | high (+2) | 63.17 | na | 30:01/68:01 | 1.38/0.99 | 08/35 | 0.95/0.82 |
| 6197-06 / 6197-04 | 03:02/07:01 | 02:02/04:02 | 02:01/04:01 | neutral (0) | 9.33 | 0.73/0.17 | 02:02/24:02 | 1/1.42 | 42/45 | 0.68 |
| 6450-02C / 6450-04A | 03:01/03:01 | 02:01/02:01 | 05:01/05:01 | moderate (+1) | 20.92 | na | 01:01/33:01 | 0.56/0.24 | 08/65 | 0.95 |
| 6429-02A / 6429-02 | 01:03/03:01 | 05:01/02:01 | 01:01/05:01 | neutral (0) | 6.96 | na/4.23 | 01:01/02:01 | 0.56/1 | 44/81 | 1 |
| 6267-02A / 6267-01 | 04:01/04:04 | 03:02/03:02 | 03:01/03:01 | high (+2) | 32.3 | na | 01:01/11:01 | 0.56/0.52 | 39/60 | 2.41 |
| 6310-02B / 6310-02B | 07:01/11:02 | 02:02/03:19 | 02:01/05:01 | neutral (0) | 1.19 | 0.5/na | 03:01/30:01 | 1.13/1.38 | 08/57 | 0.95/0.5 |
| 6209-01 / 6209-02 | 03:01/04:01 | 02:01/03:02 | 03:01/05:01 | high (+2) | 63.17 | na | 01:01/02:01 | 0.56/1 | na | na |
| 6380-04A / 6380-02 | 03:01/13:02 | 02:01/06:04 | 01:02/05:01 | moderate (+1) | 10.16 | 4.23/1.72 | 33:03/68:02 | 0.24/0.99 | 71/53 | 1.26 |
| 6371-02 / 6371-02 | 03:01/13:02 | 02:01/06:04 | 01:02/05:01 | moderate (+1) | 10.16 | na | 01:01/68:02 | 0.56/0.99 | 08/65 | 0.95 |
| 6228-02* / 6228-01* | 03:01/04:02 | 02:01/03:02 | 03:01/05:01 | high (+2) | 63.17 | na | 23:01/68:01 | 0.71/0.99 | 60/44 | 1 |
| 6396-02B / 6396-04 | 03:01/07:01 | 02:01/02:02 | 02:01/05:01 | neutral (0) | 3.24 | na | 23:01/24:02 | 0.71/1.42 | 44/49 | 1/3.23 |
| 6247-01 / 6247-03 | 04:04/11:01 | 03:01/03:02 | 03:01/05:01 | neutral (0) | 14.89 | na | 24:02/68:01 | 1.42/0.99 | 44/60 | 1 |
| 6362-04A / 6362-02 | 01:03/03:01 | 02:01/05:01 | 01:01/05:01 | neutral (0) | 6.96 | na | 03:01/11:01 | 1.13/0.52 | 18/35 | 1.83/0.82 |
| 6405-02A / 6405-04 | 03:01/04:07 | 02:01/03:02 | 05:01/03:01 | high (+2) | 63.17 | 6.95/1.20 | 30:02/31:01 | 1.38/0.94 | 18/61 | 1.83 |
| 6264-02 / 6264-01 | 03:01/04:04 | 02:01/03:02 | 03:01/05:01 | high (+2) | 63.17 | na | 23:01/32:01 | 0.71/0.37 | na | na |
| 6089-06 / 6089-03 | 01:01/04:01 | 03:02/05:01 | 01:01/03:01 | moderate (+1) | 16.14 | na | 01:01/02:01 | 0.56/1 | 60/62 | na |
| 6418-02B / 6418-01 | 04:01/11:03 | 03:02/03:01 | 03:01/05:05 | neutral (0) | 4.5 | na | 02:01/32:01 | 1/0.37 | 51/62 | 0.98 |
| 6180-02 / 6180-01 | 01:01/03:01 | 02:01/05:01 | 01:01/05:01 | neutral (0) | 6.96 | na | 01:01/02:01 | 0.56/1 | 08/27 | 0.95/0.52 |

T1D staging for AAb donors is based solely on the number of autoantibodies and in the absence of metabolic tests cannot distinguish between stage 1 and 2 T1D.

AA: African American
AMR: Admixed (Hispanic/Latino)
CEU: Caucasians of European descent

All numerical values in columns AD, AE, AG and AI represent odds ratios.

n/a: not applicable
na: not available
* insulitis assessment: the column features results from manual assessments conducted by nPOD pathologists. Our semi-automated image analyses largley reproduced these assessments yet we did not confirm insulitis cases described earlier as “low-grade” (AAb 6310; four islets with 51, 16, 14 and 14 CD45 cells, respectively) or “possibly” (T1DS 6380), nor for T1DL 6264 where insulitis was recorded in the pancreas body, a region not interrogated here. While we identified additional insulitis cases in PH sections from AAb 6424 and T1DL 6180 (c.f., Fig.5B), the only gross diagnostic discordance pertains to T1DS 6405, a severely obese donor for whom “insulitis present in all regions” is noted in the nPOD case report (visual inspection confirmed abundant CD45 cells in exocrine tissue of this donor yet few were associated with islets).

Table S3

**Table S3. Antibodies and MICSSS staining conditions.**

| Target | Clone | Species & isotype | Vendor | Staining condition | 2nd Ab staining condition | HRP-SAV |
|---|---|---|---|---|---|---|
| Proinsulin (ProINS) | GS-9A8 | Mouse (Ms) IgG1 | Novo Nordisk; also available from DHSB | 1:200<br>1h, RT | 1:800<br>30min, RT | 1:300<br>30min, RT |
| Islet amylin (IAPP) | polyclonal | Rabbit (Rb) IgG | Sigma<br>#HPA053194 | 1:1000<br>1h, RT | 1:1000<br>30min, RT | 1:300<br>30min, RT |
| Chromogranin A (CHGA) | LK2H10+PHE5 | Ms IgG1 | Novus Biologicals<br>#NBP2-342390; discontinued, available from Biocare Medical: https://biocare.net/product/chromogranin-a-antibody/ | 1:150<br>1h, RT | 1:850<br>30min, RT | 1:300<br>30min, RT |
| Glucagon (GCG) | IMD-7 | Ms IgG1 | Abcam<br>#ab82270; discontinued, available from LS Bio #LS C171152 | 1:500,<br>1h, RT | 1:850<br>30min, RT | 1:300<br>30min, RT |
| Insulin (INS) | polyclonal | Guinea pig (Gp) IgG | DAKO<br>#A0564; discontinued, now only available as ready-to-use format #IR00261-2 | 1:850<br>1h, RT | 1:1000<br>30min, RT | 1:300<br>30min, RT |
| Somatostatin (SST) | polyclonal | Rb IgG | DAKO<br>#A0566; discontinued, alternative available from Genetex #GTX60646 (Ms IgG1; clone 7G5) | 1:350<br>1h, RT | 1:750<br>30min, RT | 1:300<br>30min, RT |
| Pancreatic polypeptide (PPY) | polyclonal | Goat IgG | Novus<br>#NB100-1793 | 1:100<br>1.5h, RT | 1:250<br>30min, RT | 1:300<br>30min, RT |
| Proglucagon (ProGCG) | D16G10 | Rb IgG | Cell Signaling<br>#8233 | 1:75<br>1.5h RT | 1:300<br>30min, RT | 1:300<br>30min, RT |
| CD45 | 2B11 + PD7/26 | Ms IgG1 | DAKO<br>#M0701 | 1:100<br>2h, RT | 1:250<br>30min RT | 1:300<br>30min, RT |

Ab, antibody; RT, room temperature; HRP-SAV, horseradish peroxidase-conjugated streptavidin

### Figure S1

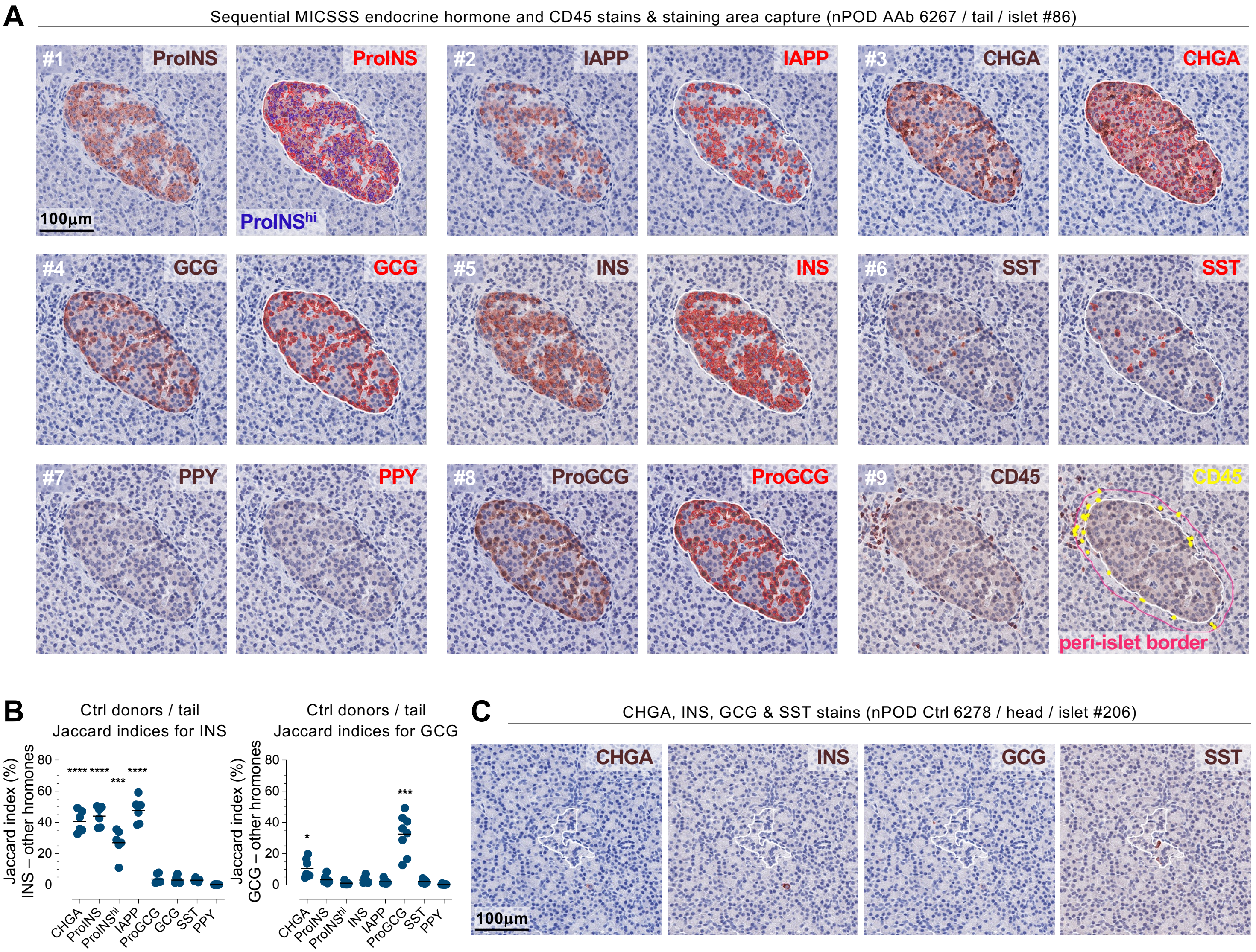
A
Sequential MICSSS endocrine hormone and CD45 stains & staining area capture (nPOD AAb 6267 / tail / islet #86)
#1 ProINS
ProINS
100μm
ProINS^hi
#2 IAPP
IAPP
#3 CHGA
CHGA
#4 GCG
GCG
#5 INS
INS
#6 SST
SST
#7 PPY
PPY
#8 ProGCG
ProGCG
#9 CD45
CD45
peri-islet border
B
Ctrl donors / tail
Jaccard indices for INS
Jaccard index (%)
INS – other hromones
0
20
40
60
80
**** **** *** ****
CHGA
ProINS
ProINS^hi
IAPP
ProGCG
GCG
SST
PPY
Ctrl donors / tail
Jaccard indices for GCG
Jaccard index (%)
GCG – other hromones
0
20
40
60
80
*
***
CHGA
ProINS
ProINS^hi
INS
IAPP
ProGCG
SST
PPY
C
CHGA, INS, GCG & SST stains (nPOD Ctrl 6278 / head / islet #206)
CHGA
INS
GCG
SST
100μm

### Figure S2

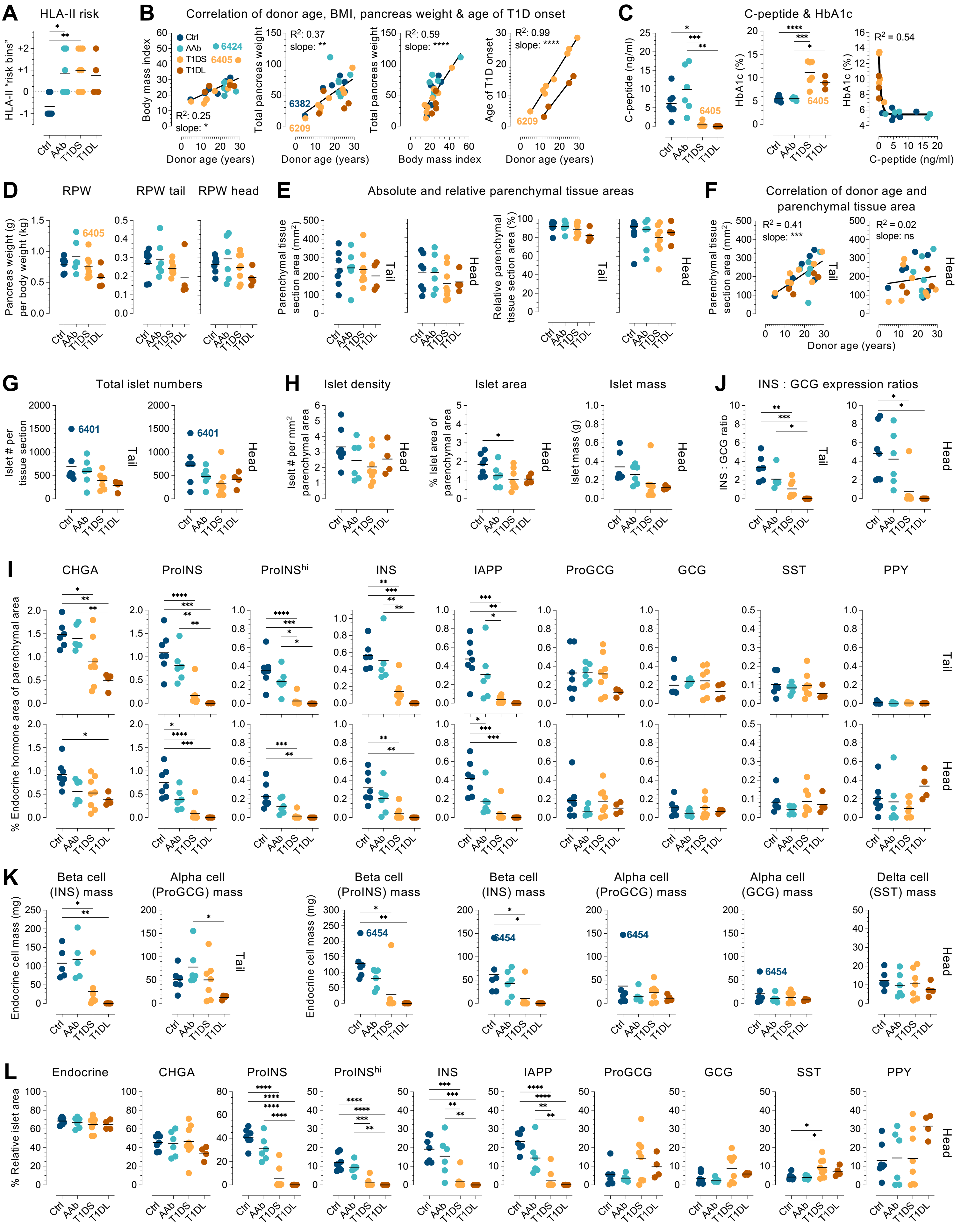

A
HLA-II risk
HLA-II "risk bins"
B
Correlation of donor age, BMI, pancreas weight & age of T1D onset
Ctrl
AAb
T1DS
T1DL
Body mass index
Total pancreas weight
Age of T1D onset
Donor age (years)
Body mass index
$R^2$: 0.25
slope: *
$R^2$: 0.37
slope: **
$R^2$: 0.59
slope: ****
$R^2$: 0.99
slope: ****
6424
6405
6382
6209
C
C-peptide & HbA1c
C-peptide (ng/ml)
HbA1c (%)
$R^2$ = 0.54
D
RPW
RPW tail
RPW head
Pancreas weight (g) per body weight (kg)
E
Absolute and relative parenchymal tissue areas
Parenchymal tissue section area (mm²)
Relative parenchymal tissue section area (%)
Tail
Head
F
Correlation of donor age and parenchymal tissue area
$R^2$ = 0.41
slope: ***
$R^2$ = 0.02
slope: ns
G
Total islet numbers
Islet # per tissue section
6401
H
Islet density
Islet area
Islet mass
Islet # per mm² parenchymal area
% Islet area of parenchymal area
Islet mass (g)
J
INS : GCG expression ratios
INS : GCG ratio
I
CHGA
ProINS
ProINS^hi
INS
IAPP
ProGCG
GCG
SST
PPY
% Endocrine hormone area of parenchymal area
K
Beta cell (INS) mass
Alpha cell (ProGCG) mass
Beta cell (ProINS) mass
Beta cell (INS) mass
Alpha cell (ProGCG) mass
Alpha cell (GCG) mass
Delta cell (SST) mass
Endocrine cell mass (mg)
6454
L
Endocrine
CHGA
ProINS
ProINS^hi
INS
IAPP
ProGCG
GCG
SST
PPY
% Relative islet area

### Figure S3

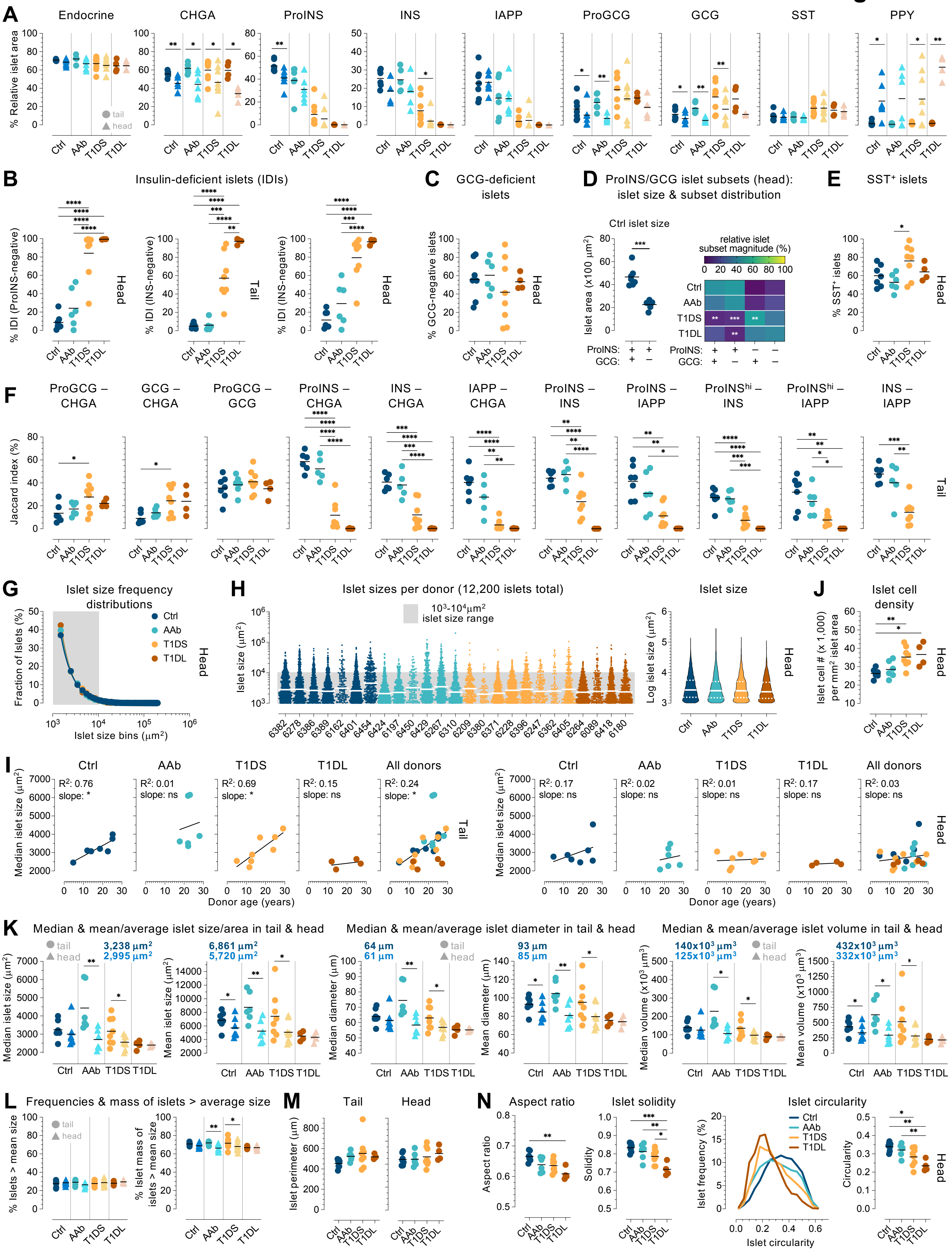
A
Endocrine
CHGA
ProINS
INS
IAPP
ProGCG
GCG
SST
PPY
% Relative islet area
tail
head
Ctrl
AAb
T1DS
T1DL
B
Insulin-deficient islets (IDIs)
% IDI (ProINS-negative)
% IDI (INS-negative)
Head
Tail
C
GCG-deficient islets
% GCG-negative islets
D
ProINS/GCG islet subsets (head): islet size & subset distribution
Ctrl islet size
Islet area (x100 μm²)
relative islet subset magnitude (%)
ProINS:
GCG:
E
SST⁺ islets
% SST⁺ islets
F
ProGCG – CHGA
GCG – CHGA
ProGCG – GCG
ProINS – CHGA
INS – CHGA
IAPP – CHGA
ProINS – INS
ProINS – IAPP
ProINShi – INS
ProINShi – IAPP
INS – IAPP
Jaccard index (%)
G
Islet size frequency distributions
Fraction of Islets (%)
Islet size bins (μm²)
H
Islet sizes per donor (12,200 islets total)
10³-10⁴μm² islet size range
Islet size (μm²)
Islet size
Log islet size (μm²)
J
Islet cell density
Islet cell # (x 1,000) per mm² islet area
I
All donors
Median islet size (μm²)
Donor age (years)
K
Median & mean/average islet size/area in tail & head
Median & mean/average islet diameter in tail & head
Median & mean/average islet volume in tail & head
Median islet size (μm²)
Mean islet size (μm²)
Median diameter (μm)
Mean diameter (μm)
Median volume (x10³ μm³)
Mean volume (x10³ μm³)
L
Frequencies & mass of islets > average size
% Islets > mean size
% Islet mass of islets > mean size
M
Tail
Head
Islet perimeter (μm)
N
Aspect ratio
Islet solidity
Islet circularity
Solidity
Islet frequency (%)
Circularity

### Figure S4

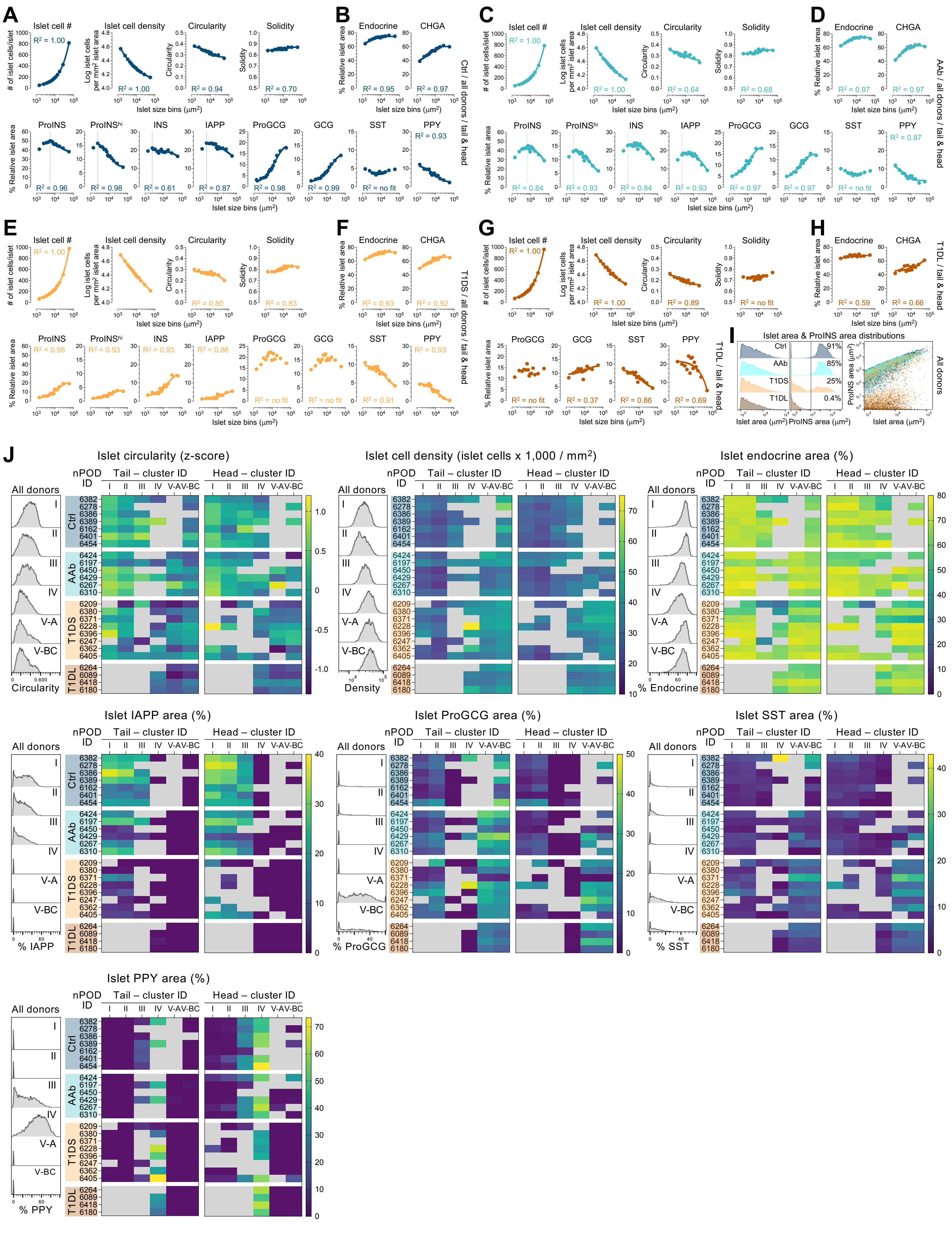

A
Islet cell #
Islet cell density
Circularity
Solidity
B
Endocrine
CHGA
Ctrl / all donors / tail & head
C
D
AAb / all donors / tail & head
E
F
T1DS / all donors / tail & head
G
H
T1DL / tail & head
I
Islet area & ProINS area distributions
All donors
J
Islet circularity (z-score)
Islet cell density (islet cells x 1,000 / mm²)
Islet endocrine area (%)
Islet IAPP area (%)
Islet ProGCG area (%)
Islet SST area (%)
Islet PPY area (%)

### Figure S5A/B

Figure S5A/B

Pancreatic tail – islet properties across UMAP clusters grouped according to T1D stage

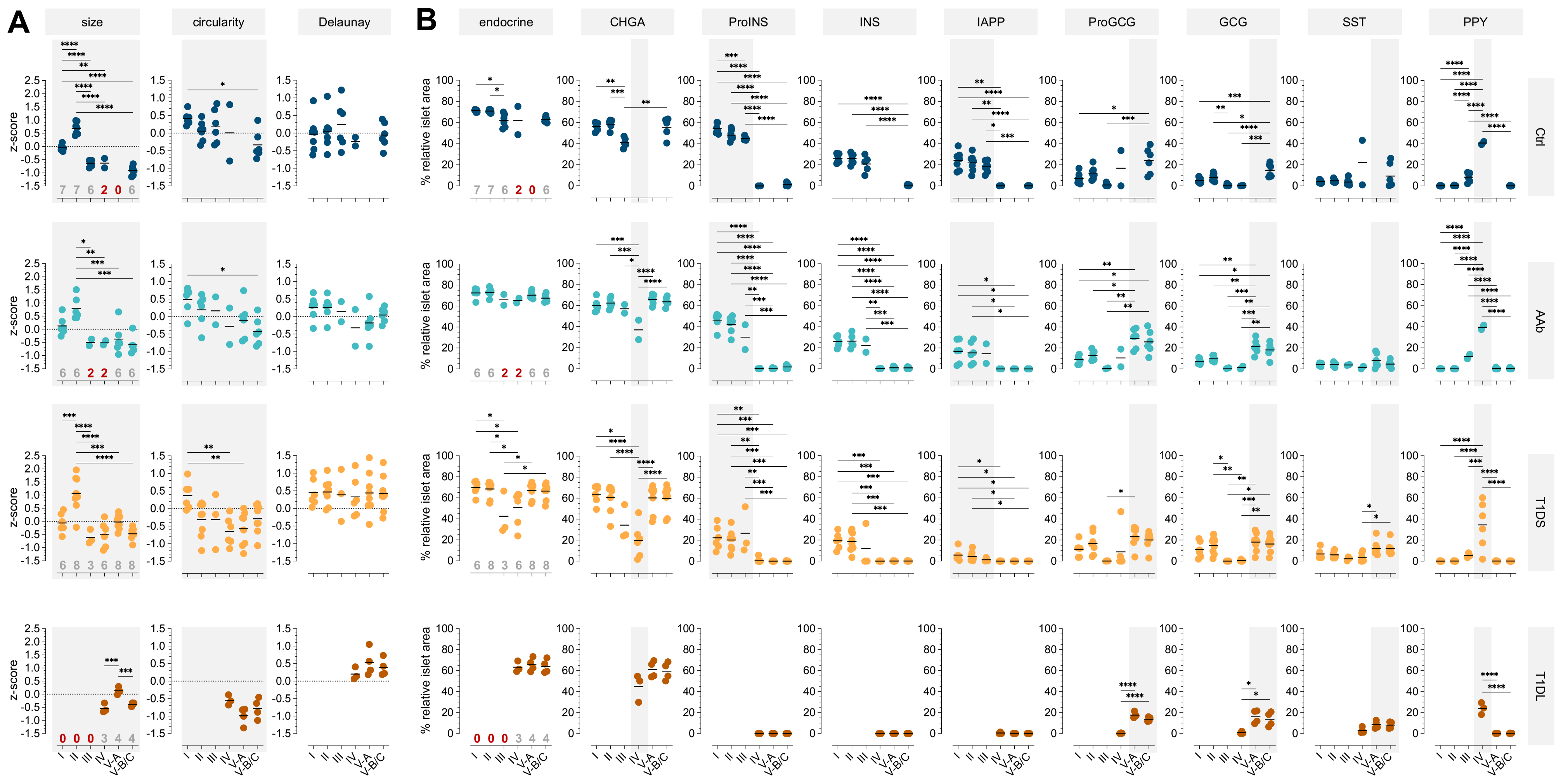

### Figure S5C/D

Figure S5C/D

Pancreatic head – islet properties across UMAP clusters grouped according to T1D stage

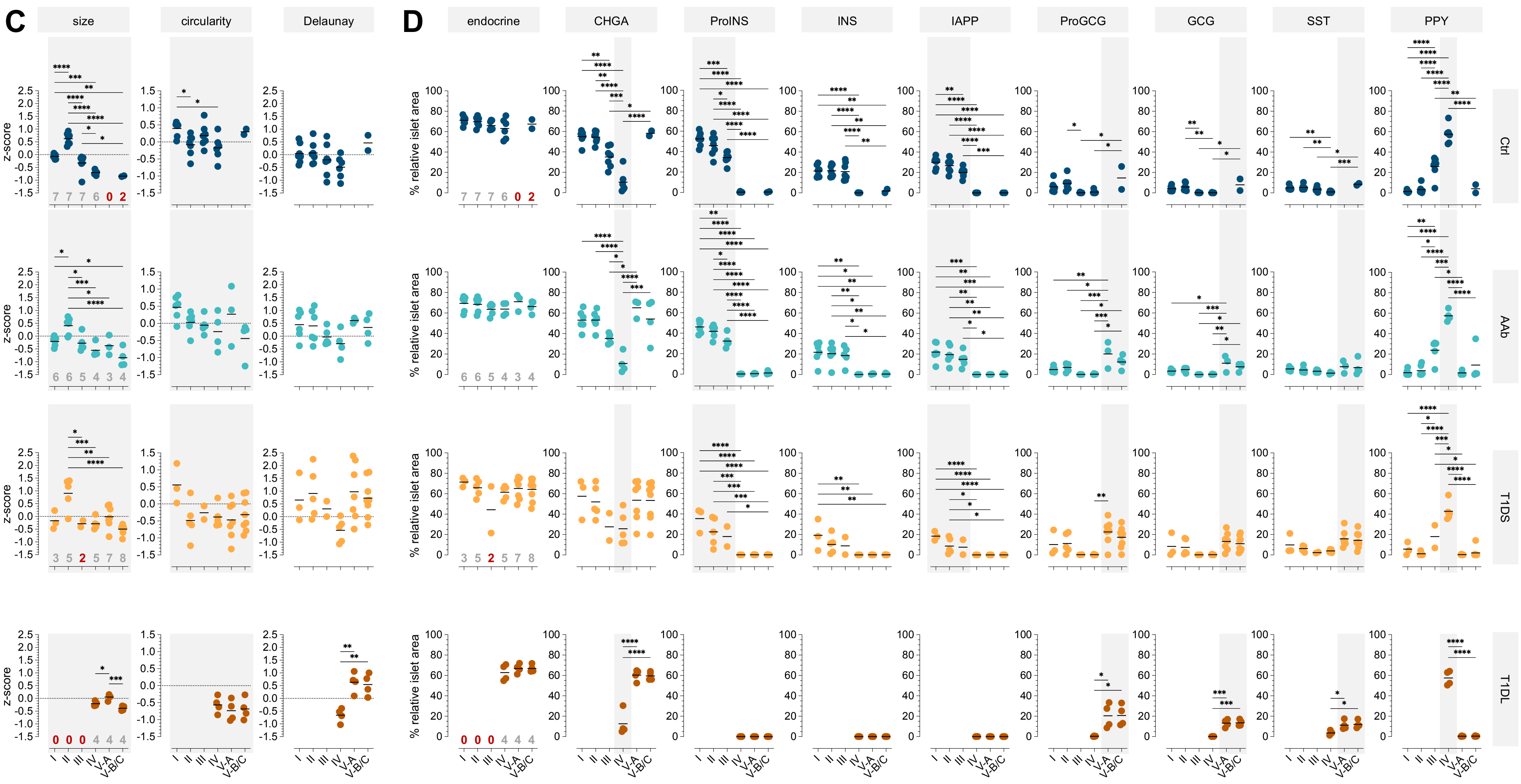

### Figure S5E/F

Figure S5E/F

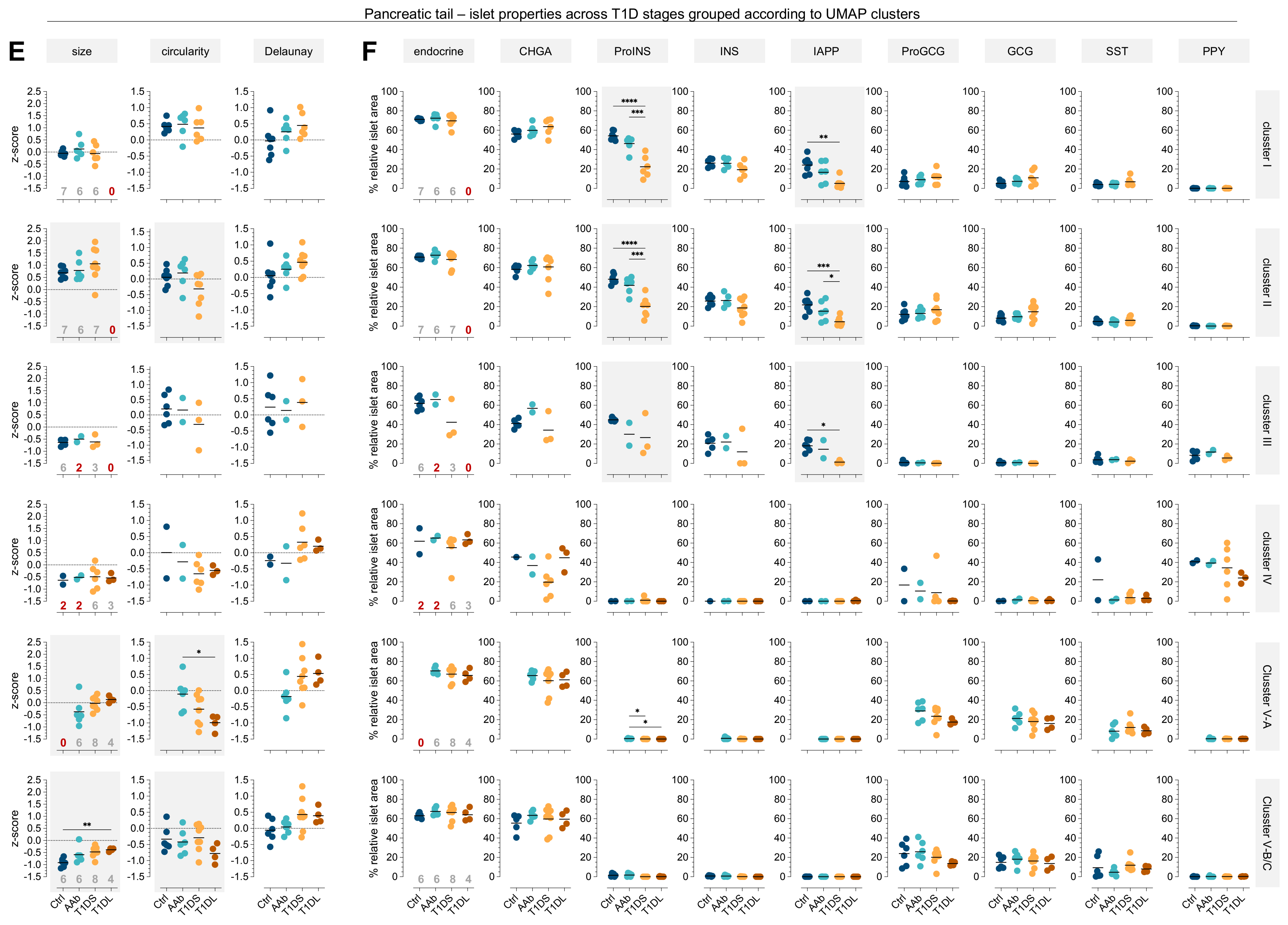

Figure S5G/H

Figure S5G/H

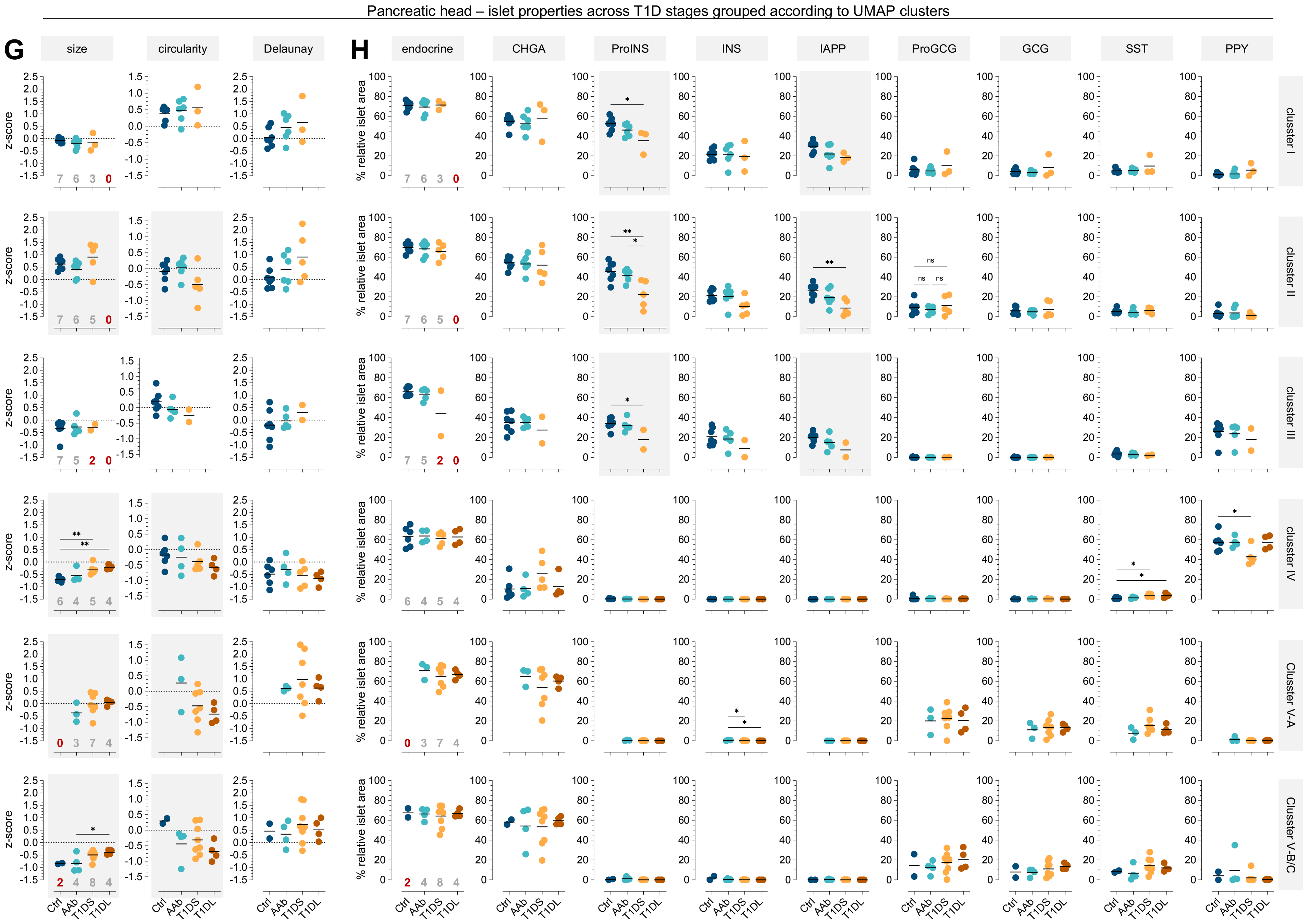

### Figure S6

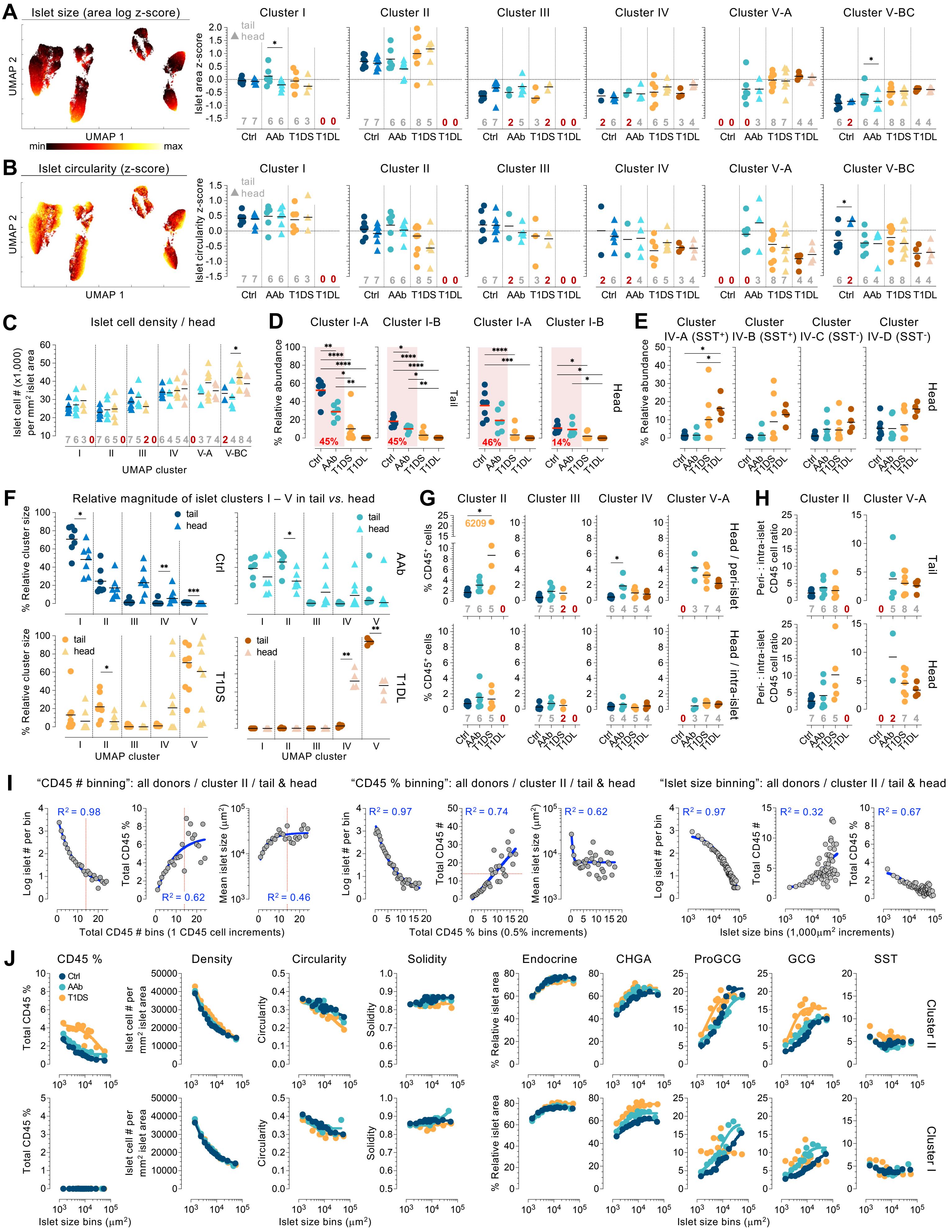

A
Islet size (area log z-score)
UMAP 2
UMAP 1
min
max
Cluster I
Cluster II
Cluster III
Cluster IV
Cluster V-A
Cluster V-BC
tail
head
Islet area z-score
Ctrl
AAb
T1DS
T1DL
B
Islet circularity (z-score)
Islet circularity z-score
C
Islet cell density / head
Islet cell # (x1,000) per mm² islet area
UMAP cluster
D
Cluster I-A
Cluster I-B
% Relative abundance
Tail
Head
45%
45%
46%
14%
E
Cluster IV-A (SST⁺)
Cluster IV-B (SST⁺)
Cluster IV-C (SST⁻)
Cluster IV-D (SST⁻)
F
Relative magnitude of islet clusters I – V in tail vs. head
% Relative cluster size
G
Cluster II
Cluster III
Cluster IV
Cluster V-A
% CD45⁺ cells
6209
Head / peri-islet
Head / intra-islet
H
Peri- : intra-islet CD45 cell ratio
I
"CD45 # binning": all donors / cluster II / tail & head
"CD45 % binning": all donors / cluster II / tail & head
"Islet size binning": all donors / cluster II / tail & head
Log islet # per bin
Total CD45 %
Mean islet size (µm²)
Total CD45 #
R² = 0.98
R² = 0.62
R² = 0.46
R² = 0.97
R² = 0.74
R² = 0.62
R² = 0.97
R² = 0.32
R² = 0.67
Total CD45 # bins (1 CD45 cell increments)
Total CD45 % bins (0.5% increments)
Islet size bins (1,000µm² increments)
J
CD45 %
Density
Circularity
Solidity
Endocrine
CHGA
ProGCG
GCG
SST
Islet cell # per mm² islet area
% Relative islet area
Cluster II
Cluster I
Islet size bins (µm²)

### Figure S7

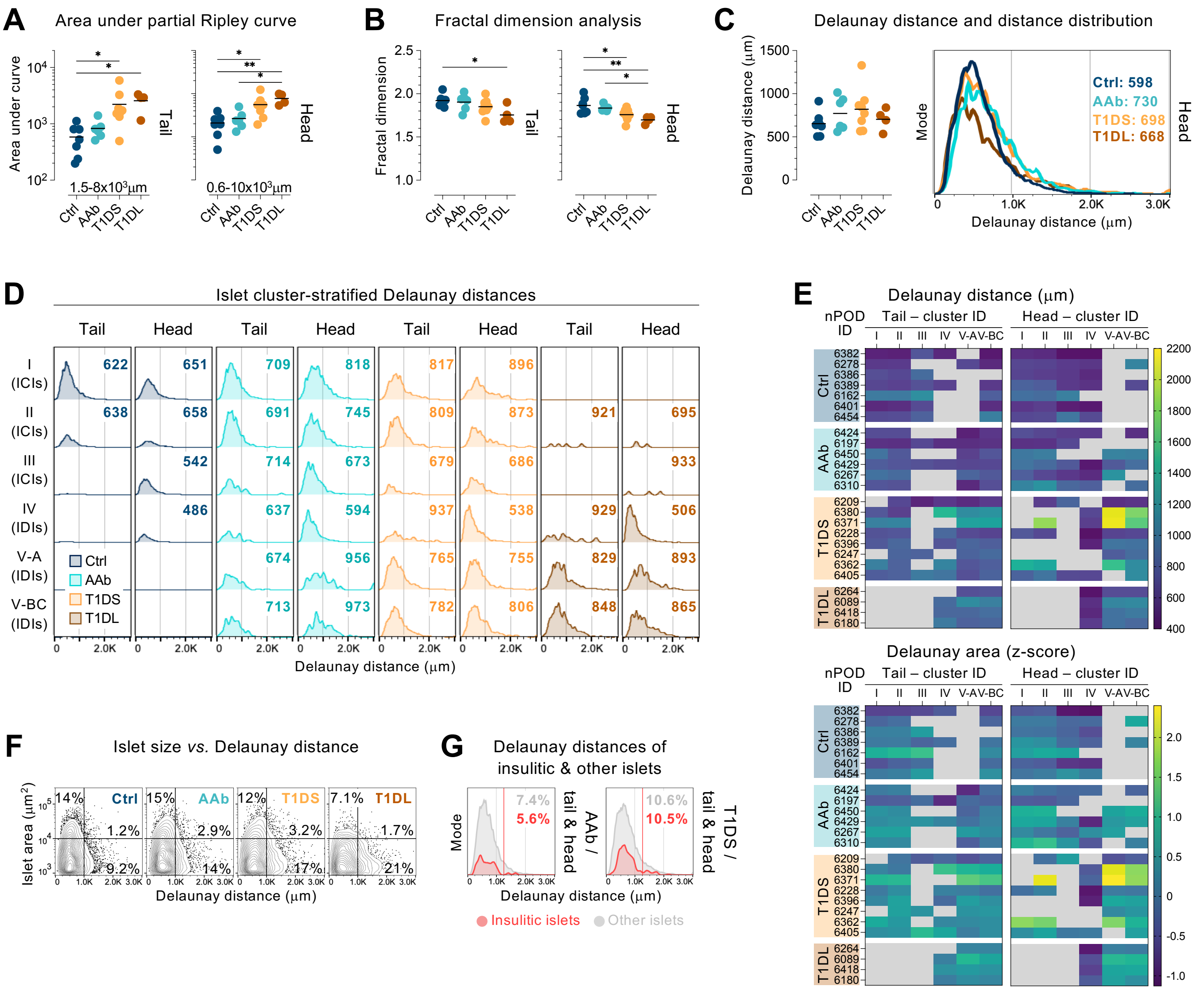


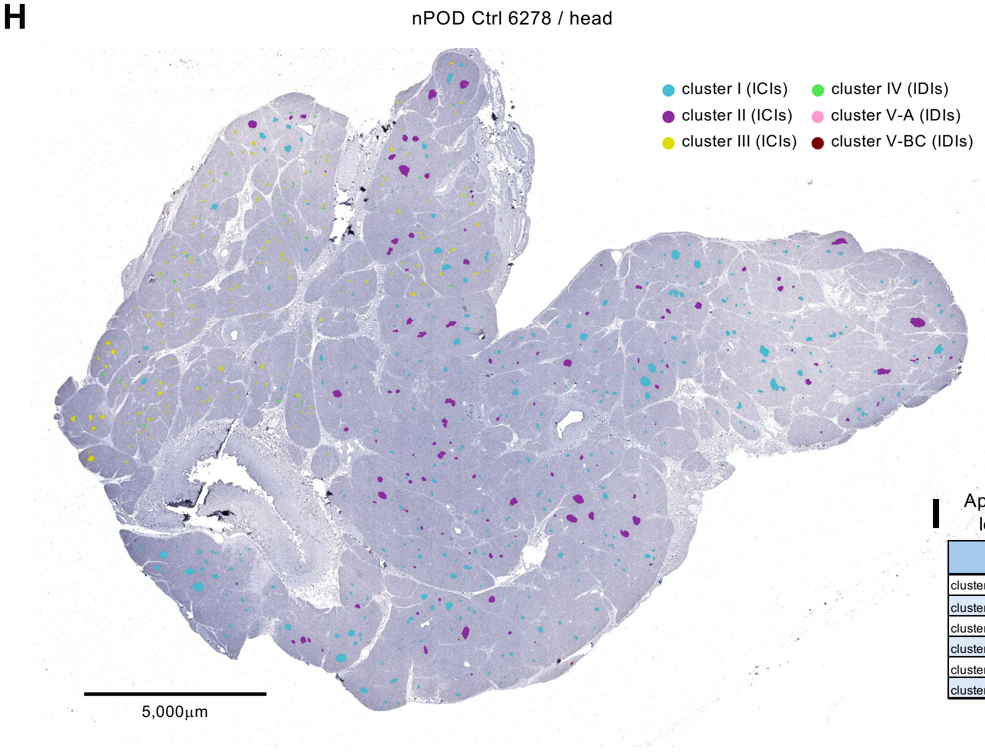


**I** Approximation of "islet clustering" with low dimensionality immunostaining

| | INS | GCG | peri- & intra islet CD45 | PPY |
|---|---|---|---|---|
| cluster I | + | + or - | 0 | - |
| cluster II | + | + or - | ≥ 1 | - |
| cluster III | + | - | ≥ 0 | + |
| cluster IV | - | - | ≥ 0 | + |
| cluster V-A | - | + | ≥ 1 | - |
| cluster V-BC | - | + | 0 | - |

### Figure S8

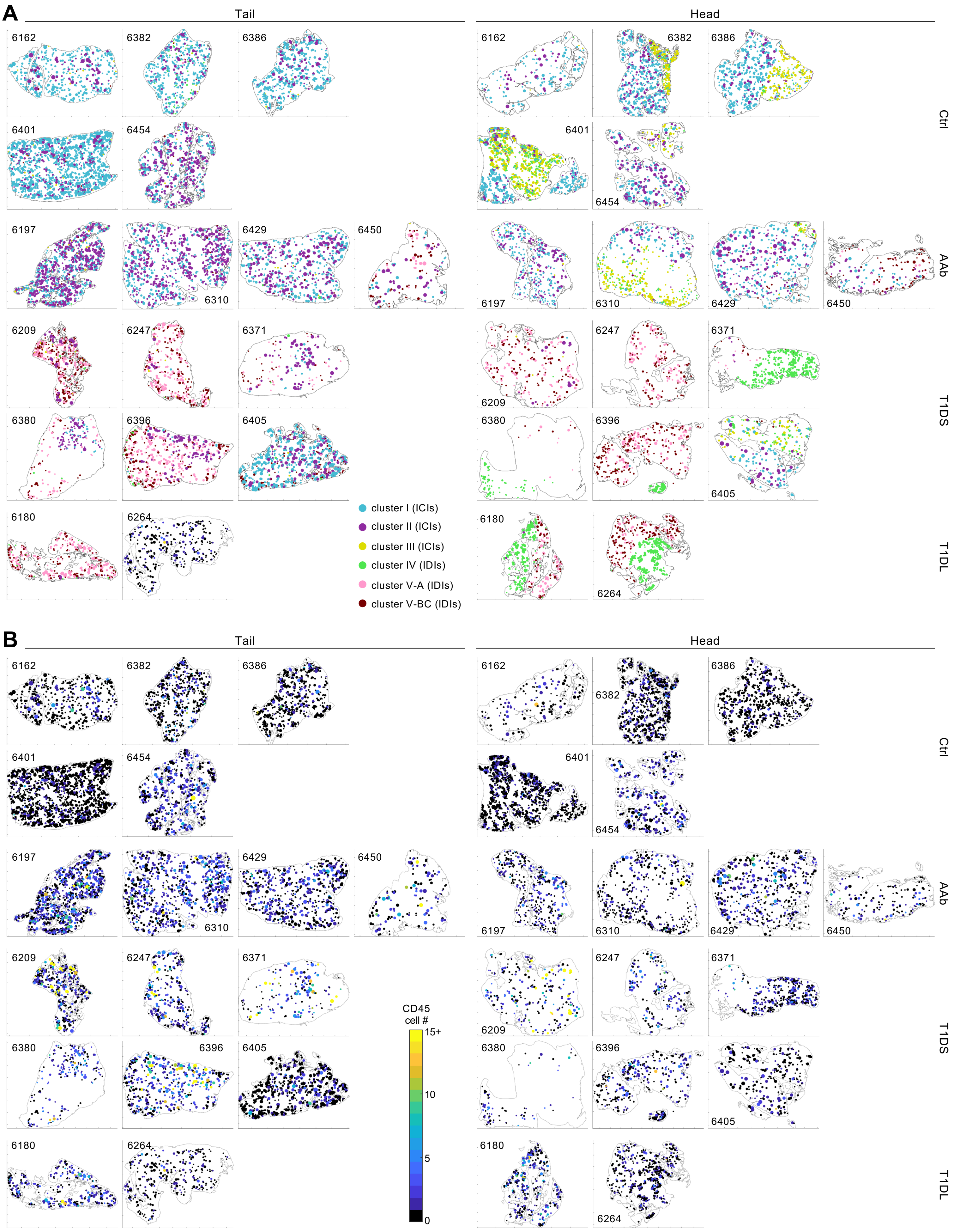
A
Tail
Head
6162
6382
6386
6401
6454
Ctrl
6197
6310
6429
6450
AAb
6209
6247
6371
6380
6396
6405
T1DS
6180
6264
T1DL
cluster I (ICIs)
cluster II (ICIs)
cluster III (ICIs)
cluster IV (IDIs)
cluster V-A (IDIs)
cluster V-BC (IDIs)
B
Tail
Head
CD45
cell #
15+
10
5
0
